# Mechanisms of HIV-1 Integrase Resistance to Dolutegravir and Potent Inhibition of Drug Resistant Variants

**DOI:** 10.1101/2022.12.04.519057

**Authors:** Min Li, Dario Oliveira Passos, Zelin Shan, Steven J. Smith, Qinfang Sun, Avik Biswas, Indrani Choudhuri, Timothy S. Strutzenberg, Allan Haldane, Nanjie Deng, Zhaoyang Li, Xue Zhi Zhao, Terrence R. Burke, Ronald M. Levy, Stephen H. Hughes, Robert Craigie, Dmitry Lyumkis

## Abstract

HIV-1 infection depends on the integration of viral DNA into host chromatin. Integration is mediated by the viral enzyme integrase and is blocked by integrase strand transfer inhibitors (INSTIs), first-line antiretroviral therapeutics widely used in the clinic. Resistance to even the best INSTIs is a problem and the mechanisms of resistance are poorly understood. Here, we analyze combinations of the mutations E138K, G140A/S, and Q148H/K/R, which confer resistance to INSTIs. The investigational drug 4d more effectively inhibited the mutants compared with the approved drug Dolutegravir (**DTG**). We present 11 new cryo-EM structures of drug resistant HIV-1 intasomes bound to **DTG** or **4d**, with better than 3 Å resolution. These structures, complemented with free energy simulations, virology, and enzymology, explain the mechanisms of **DTG** resistance involving E138K+G140A/S+Q148H/K/R and show why **4d** maintains potency better than **DTG**. These data establish a foundation for further development of INSTIs that potently inhibit resistant forms in integrase.

## INTRODUCTION

HIV-1 and other retroviruses integrate a DNA copy of their viral RNA genome into the genome of the host cell, a process that is mediated by the virally encoded enzyme integrase (IN) (*1, 2*). The integrated proviruses serve as templates for transcription of new viral RNA genomes and the mRNAs required for production of progeny virions. The catalytic integration reaction that inserts a DNA copy of the viral genome in the host genome presents a target for treating people living with HIV-1 (PLWH) and for preventing new infections.

Two chemical steps are required for integration: (*i*) 3’-end processing, in which IN cleaves two nucleotides from each 3’-end of the viral DNA, and (*ii*) DNA strand transfer, in which IN inserts the ends of the viral DNA into target DNA in a one-step transesterification reaction (*3*). Completion of integration requires removal of two nucleotides from the 5’-ends of the viral DNA, filling in of the single strand gaps between viral and target DNA, and ligation. These latter steps are mediated by cellular enzymes. IN is both necessary and sufficient to carry out both 3’-processing and DNA strand transfer *in vitro*. Each step of DNA integration, up to formation of the integration intermediate, occurs within a series of stable nucleoprotein complexes (intasomes), involving a multimer of IN and a pair of viral DNA ends. The atomic structures have now been determined for a variety of retroviral intasomes (*4–13*), including HIV-1 (*14, 15*), revealing differences in the oligomeric assemblies and how they engage target DNA. The formation of intasome species, and the intasome-mediated insertion of the vDNA into the host genome, are essential steps in the viral replication cycle [see (*1, 2*) for recent reviews].

Because integration is an essential step in viral replication, intasomes are targeted by an important class of antiretroviral drugs, the IN strand transfer inhibitors (INSTIs). INSTIs selectively interact with both the bound viral DNA and IN and, as a consequence, do not bind tightly to free IN (*4, 16, 17*). INSTIs work by chelating the two Mg^2+^ ions, blocking the active site and inhibiting the strand transfer activity of IN. Five INSTIs have been clinically approved by the US Food and Drug Administration and antiretroviral regimens that include INSTIs are standard of care. These include 1^st^ generation INSTIs Raltegravir (**RAL**, approved 2007) and Elvitegravir (**EVG**, approved 2011), as well as 2^nd^ generation INSTIs Dolutegravir (**DTG**, approved 2013), Bictegravir (**BIC**, approved 2018) and Cabotegravir (**CAB**, approved 2021). INSTIs are widely considered to be among the best therapeutic options for use in combination therapies to treat HIV-infected patients (*13*).

The major difference between 1^st^ and 2^nd^ generation INSTIs lies in their ability to inhibit mutant forms of IN. Whereas both **RAL** and **EVG** can potently inhibit the WT enzyme, drug resistant mutations (DRMs) quickly develop (*19*). The 2^nd^ generation INSTIs can potently inhibit many of the mutant forms of IN that arise during the course of treatment with the 1^st^ generation inhibitors (*20–22*). However, clinical data indicates that resistance to 2^nd^ generation INSTIs is now a growing problem (*23–25*). Unfortunately, because HIV-1 replicates rapidly and the viral load is high, drug-resistant mutations can arise, and viral sequences within an infected individual can differ by 10% or more (*26*). These data highlight the challenges to developing effective treatments. They also point to the fact that, while resistance is a problem, it can be addressed when the mechanisms are understood and by using this information to develop compounds that inhibit both the WT and the DRM enzymes.

The most frequently encountered DRMs to INSTI therapy include mutations at position Q148H/K/R. The Q148 mutations are commonly found in combination with the G140A/S mutations (*23–25, 27*) [and, rarely, G140C (*27*)]. In addition to the pair of DRMs at positions 140/148, numerous other DRMs arise in and around the active site, frequently including E138K (*27*). Recent structural work using the Simian Immunodeficiency Virus (SIVrcm) IN as a model for HIV-1 IN, led to a proposed mechanism for the resistance of the G140S/Q148H double mutant (*28*). The mutation Q148H arises first, expelling a water molecule from the secondary coordinating shell of the Mg^2+^ ions and introducing a bulky electropositive residue underneath the bound ligand. The subsequent mutation of the neighboring residue G140S acidifies the Nε2 of H148. Coupling of the residues S140-H148-E152 increases the partial positive charge near the Mg^2+^ ligand. Collectively, these changes weaken chelation of the bound INSTI, leading to a reduction in drug binding. Because there are many complex DRMs that have a change at Q148 (H/K/R), Mg^2+^ chelation of Mg^2+^ was suggested to be the Achilles’ heel of this drug class. However, SIVrcm serves as a proxy for HIV-1; their active sites, including residues that are encountered in drug-resistant viral clones, are not identical. Moreover, only the double mutant G140S/Q148H was structurally defined, but there are many combinations of mutations at position 148H/K/R which can be paired with the G140A/S mutations and with other nearby mutations, including E138K. Thus, the full consequences of these and related mutations, and the structural basis of resistance in HIV-1 IN, which should help us understand how to develop INSTIs that can overcome resistance caused by mutations at these positions, remains unclear.

INSTIs containing a naphthyridine core have been developed into potent IN inhibitors that rival or even outperform clinical drugs in terms of their ability to inhibit a broad range of DRMs (*20, 21, 29, 30*). For example, the compound **4d** was reported to inhibit certain DRMs, including variants with 140/148 mutations, with over an order of magnitude higher potency than the clinically used drug **DTG** (*21*). Structural biology data has shed light on how naphthyridine-containing compounds bind to intasomes from HIV-1 and Prototype Foamy Virus (PFV) (*15, 29*), revealing insights into the small but significant differences in the ways INSTIs bind to PFV and HIV-1 IN (*15*). However, how and why these compounds retain potency against DRMs remains unclear. Thus, there is a pressing need to explain not only the underlying mechanisms of viral resistance, but also to understand why compounds like **4d** can potently inhibit resistant forms of IN.

Defining the underlying mechanisms of resistance lags behind clinical tabulation of emerging variants. This is primarily due to a lack of high-resolution structures of resistant variants and complementary tools to probe the resistance mechanisms. There have also been discrepancies in resistance profiles recorded for the 140/148 positions, and their behavior in combination with other DRMs. Since mutations in HIV-1 IN that confer drug resistance occur in several distinct combinations, structures of HIV-1 intasomes bound to inhibitors with and without such mutations, complemented by insights on inhibitory potencies and enzyme/viral fitness, are required to understand the details of how these mutations confer resistance and to guide the design of drugs with improved resistance profiles.

Here, we sought to understand the mechanism(s) of resistance caused by mutations at positions 138, 140, and 148. Our results reveal weaknesses in one of the leading clinically used drugs, **DTG** and explain the mechanism(s) that underly its loss of potency against DRMs. We extend these findings to explain how and why mutations at the three residue substitutions lead to drug resistance. Lastly, we provide insight into how a leading investigational compound, which is currently undergoing pre-clinical testing, retains considerable potency against the set of **DTG**-resistant variants we analyzed. Collectively, these data provide mechanistic insights into both drug resistance and its subversion and will enable more direct pathways to designing INSTIs with improved resistance profiles.

## RESULTS

### Inhibitory potencies of DTG and a leading developmental INSTI 4d against resistant mutants

Novel developmental INSTIs based on the naphthyridine scaffold maintain potency against drug resistant variants with mutations at position Q148 of HIV-1 IN (*15, 20, 21, 29–31*). However, whether their potency is maintained across the Q148H/K/R family of DRMs, particularly in association with the E138K and G140A/S mutations, remains unclear. We determined the ability of a leading clinically used 2^nd^ generation INSTI, **DTG,** to inhibit mutant viruses containing all possible combinations of E138K+G140A/S+Q148H/K/R DRMs using single-round viral infection assays and compared the data to the most potent pre-clinical developmental INSTI **4d** (**Fig. 1A**).

**Figure 1:**
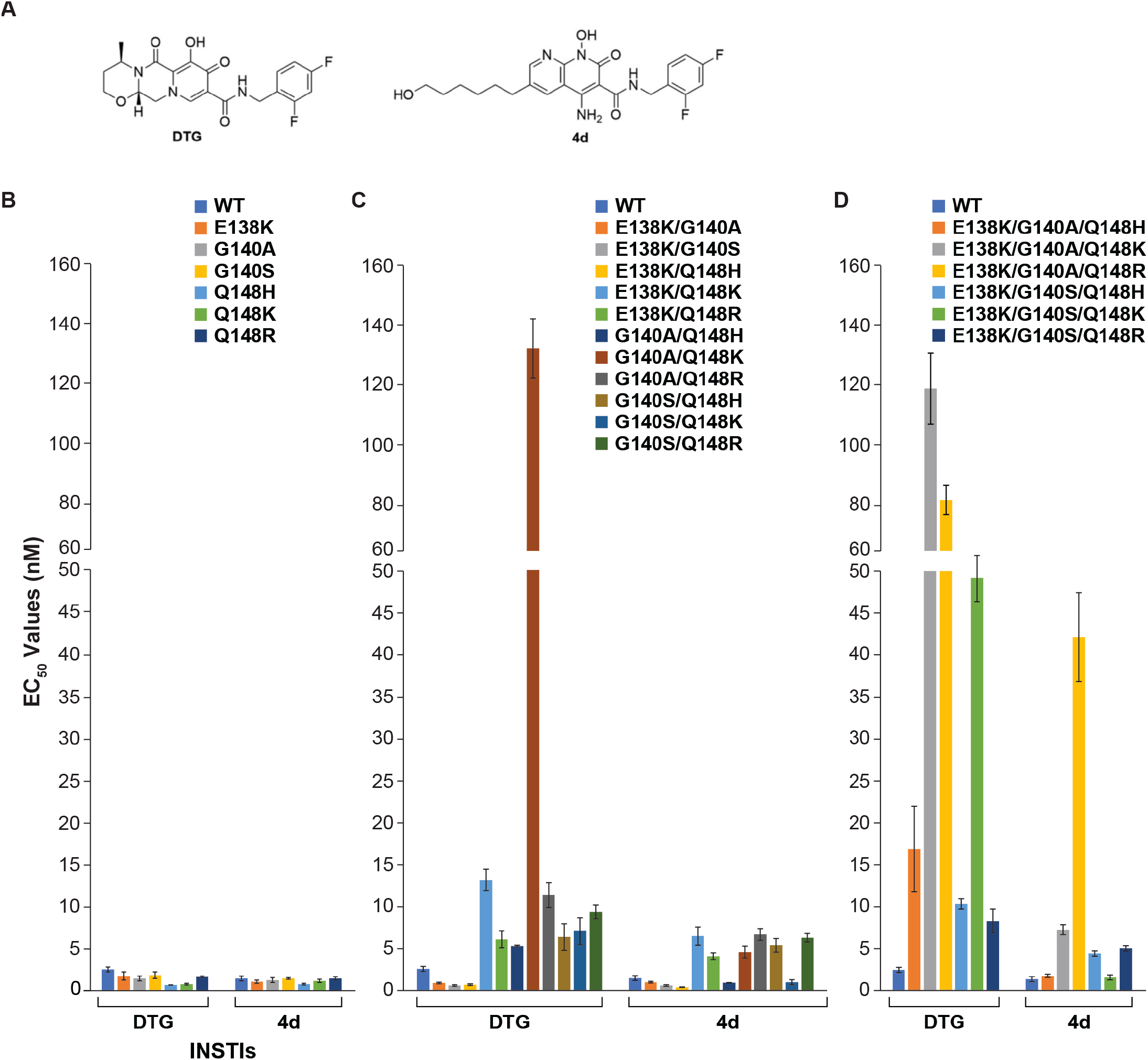
Comparison of antiviral potencies of DTG and 4d against a panel of INSTI-resistant mutants. (**A**) Chemical structures of **DTG** and **4d**, used for the assays. The EC_50_ values were determined for either of the two compounds using vectors that carry INSTI-resistant (**B**) single, (**C**) double, and (**D**) triple mutants in a single-round infection assay. The error bars represent standard deviations of independent experiments, n=3, performed in triplicate.

We began by measuring the EC_50_ values against the single mutants in a single-round infection assays, which include E138K, G140A, G140S, Q148H, Q148K, and Q148R (**Fig. 1B**). Both **DTG** and **4d** potently inhibited WT HIV-1 with EC_50_ values of 2.6 ± 0.3 nM and 1.5 ± 0.3 nM, respectively. The compounds also retained full potency (<2.0 nM) against all the mutants E138K, G140A/S, Q148H/K/R.

We next tested the IN double mutants E138K/G140A, E138K/G140S, E138K/Q148H, E138K/Q148K, E138K/Q148R, G140A/Q148H, G140A/Q148K, G140A/Q148R, G140S/Q148H, G140S/Q148K, and G140S/Q148R (**Fig. 1C**). To compare the relative effectiveness of the two drugs, we calculated the fold changes (FCs) in the EC_50_ values, using the Monogram biological cutoffs for resistance for **DTG** as a guide (*32*). Generally, both **DTG** and **4d** retained potency against this panel of IN double mutants, but **4d** has a moderately better antiviral profile compared to **DTG**. **4d** potently inhibited the IN double mutant G140A/Q148K (4.6 ± 0.7 nM; FC = 3), whereas **DTG** did not (132.0 ± 9.9 nM; FC = 51, p-value <0.001). This suggests that the G140A/Q148K mutant has specific properties that make it particularly susceptible to **4d** but not to **DTG.**

Last, we tested the complex IN mutants containing three or more mutations, which include E138K/G140A/Q148H, E138K/G140A/Q148K, E138K/G140A/Q148R, E138K/G140S/Q148H, E138K/G140S/Q148K, and E138K/G140S/Q148R (**Fig. 1D**). **4d** was considerably more effective than **DTG** against IN mutants E138K/G140A/Q148K (7.3 ± 0.6 nM; FC = 5 and 117.6 ± 11.6 nM; FC = 45, p-values <0.001, respectively), and E138K/G140S/Q148K (1.7 ± 0.3 nM; FC = 1 and 49.2 ± 2.7 nM; FC = 19, p-values <0.001, respectively). These results substantiate our observation that the double mutant G140A/Q148K is resistant to **DTG** and suggest that E138K can, under some circumstances, potentiate resistance. In accordance with the latter observation, the IN triple mutants formed by the addition of E138K to the IN double mutants G140A/Q148H, G140S/Q148H, G140S/Q148K, and G140A/Q148R all showed a decrease in susceptibility to **DTG**, although G140A/Q148K and G140S/Q148R remained largely unchanged. The only one of these triple mutants that led to a loss of susceptibility to **4d** inhibition was E138K/G140A/Q148R. Although **4d** did lose potency against E138K/G140A/Q148R (41.5 ± 5.2 nM; FC = 28), it retained more potency than **DTG** (p-value <0.001). Overall, as with the double mutants, the antiviral profile of **4d** was always better than **DTG** against this panel of triple mutants.

Collectively, we find that there are significant variations in potency among the Q148H/K/R variants, but combinations containing Q148K were consistently less susceptible to **DTG** inhibition. Mutations at position G140A/S typically potentiate the resistance of Q148H/K/R, especially variants containing Q148K. The mutation E138K can potentiate resistance in the Q148H/K/R series, sometimes decreasing the potency of an INSTI by up to an order of magnitude, although this effect was less consistent in comparison to G140A/S. Importantly, the pre-clinical INSTI **4d** retained potency better than **DTG** against all tested mutants. This difference is most apparent in the ~10-20-fold higher potency against the double mutant G140A/Q148K and the triple mutant E138K/G140A/Q148K.

### Fitness of the mutant viruses

DRMs that arise from the effects of drug pressure can have pleiotropic effects on the fitness of mutant viruses, which affect viral replicative capacity and derive from numerous factors, including protein stability and *in vitro* integration activity. Because the effects of the mutations on viral fitness affect which mutations are selected in vivo, we wanted to understand the effects of introducing DRMs on the fitness of the mutant viruses.

We first tested the ability of mutant INs to perform two-ended integration *in vitro* using double-stranded oligonucleotide mimics of the U5 vDNA end in a strand transfer assay (**Fig. 2A-B**). In comparison to the WT enzyme, mutations Q148H/K/R reduced integration activity of IN ~5-10-fold, and mutations at G140A/S reduced integration activity ~3-5-fold. When G140A/S mutations were paired with Q148H/K/R mutations, the reduction in integration typically matched the lowest integration activity for the single mutants, with the sole exception being the double mutant pair G140S/Q148H; when paired with Q148H, but not with Q148K/R, the G140S mutation restored integration by ~3-5-fold. We also noted the compensatory behavior of E138K. By itself, this mutation has a minimal effect on integration, but when other changes are present, E138K consistently restores integration activity (e.g., compare Q148K with E138K/Q148K or G140A/Q148K with E138K/G140A/Q148K). The compensatory behavior of E138K is observed across experiments, leading to an increase in the integration activity of all of the triple mutant INs in comparison to their double or single mutant counterparts.

**Figure 2:**
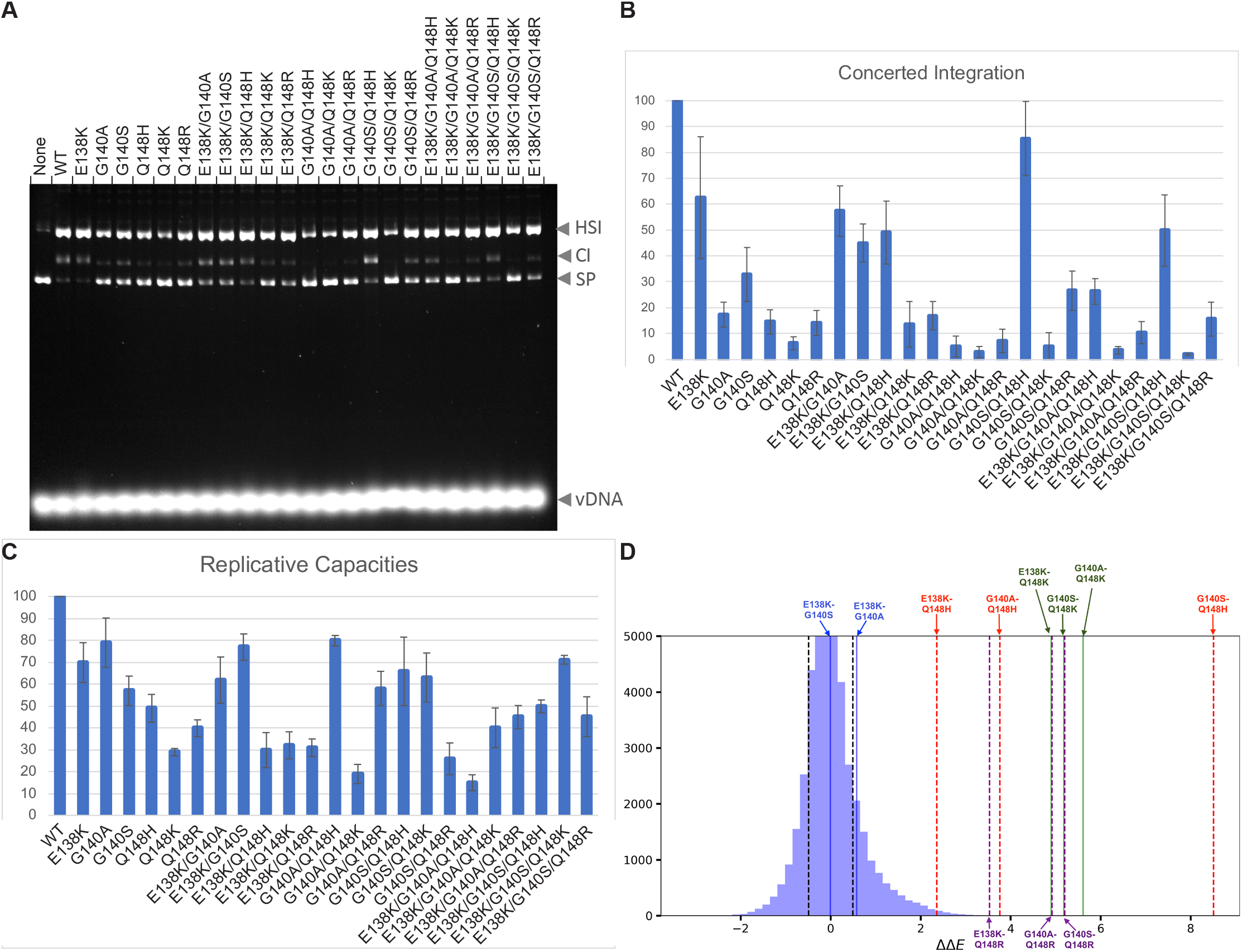
Fitness measurements. (**A**) Enzymatic catalytic activities, half-site (HS) and concerted integration (CI), for all combinations of single, double, and triple IN mutants corresponding to changes at positions E138K+G140A/S+Q148H/K/R. The bands corresponding to supercoiled plasmid DNA (SP) and viral DNA are also indicated. (**B**) Bar graph of the two-end integration data in B. (**C**) Replicative capacities measured in cell culture of the INSTI-resistant mutants using a single-round infection assay. (**D**) Double mutant cycle data derived from Potts statistical analyses. As fitness is inversely proportional to the Potts energy, ΔΔE > 0 implies that the mutations are compensatory. The plot shows that many of the residue pairs, most notably G140S+Q148H and G140A+Q148K, are coupled to one another, indicating that compensatory effects between these pairs of positions within the protein drive the observed residue co-variation. Residue pairs close to the center of the distribution are not coupled.

To determine the relative fitness of WT HIV-1 or the IN single, double, and triple mutant viruses in the absence of drugs, we used a single round infection assay (**Fig. 2C**). Among the IN single mutants, the relative infectivities of E138K, G140A, and G140S were the highest, whereas Q148H/K/R mutants reduced the replicative capacities to greater extents. Our data reproduces prior reports indicating that G140S acts as a compensatory mutation in the context of Q148H (*28, 33*), and we additionally found a modest (~2-fold) compensation for loss of replicative capacity when G140S was added to Q148K and when G140A was added to Q148R. The addition of E138K as a third mutation to the IN double mutants G140A/S + Q148H/K/R had varying effects in single round replication assays. Whereas the addition of E138K played a compensatory role for the IN triple mutants E138K/G140S/Q148K and E138K/G140A/Q148K, its effect in the context of other mutants was variable. These data suggest that the G140S mutation can have, in some cases, a compensatory effect when combined with a Q148H/K mutation and the E138K mutation can also have, in some cases, a compensatory effect when combined with the G140A/S + Q148H/K/R mutations.

Generally, we observed a similar decrease in integration activity in mutant INs when the values are compared with replicative capacities of mutant viruses, although these different measures of the effects on IN mutations did not exactly align for all of the DRMs. When the values do not align, we typically observed a decrease in integration, but not replicative capacity. It is possible that there is a modest excess of IN enzymatic activity in the virion, so that a small loss of enzymatic activity would not have a significant effect in an infectivity assay. In addition, mutations in IN can quantitatively affect the efficiency of *in vitro* reactions by mechanisms that might not be relevant to integration in cells. For example, IN tends to aggregate, and the intasome forms non-physiologically relevant stacks under *in vitro* reaction conditions (*14, 34*). Both the propensity of IN to aggregate and form stacks is exquisitely sensitive to reaction conditions and mutations in the IN protein. Thus, the readout from *in vitro* integration assays does not necessarily quantitatively mirror the intrinsic enzyme activity of IN, which would, in turn, explain the unusually low integration activity in some cases.

We next extended the *in vitro* measurements, conducted with either purified proteins or in cell culture, to patient sequence data. To this end, we took advantage of the vast amount of IN sequences derived from drug-experienced patients available in the Stanford HIV Drug Resistance Database (*35, 36*). We use a co-evolutionary (Potts) Hamiltonian model to define the underlying relationships between protein structure, function, and fitness. The Potts Hamiltonian provides a mapping of sequences to statistical scores in which lower Potts statistical energies are more probable, generating a “prevalence landscape” (*37, 38*). Recent studies have demonstrated that experimental measurements of fitness differences are well correlated with the change in Potts statistical energy for sequences from large Eukaryotic protein families (*39, 40*). Potts models have also been used to interrogate the fitness landscape of viral systems, most notably HIV-1 proteins escaping immune pressure (*37, 41*) and HIV-1 enzymes evolving under drug selection pressure (*42, 43*).

Here, we utilized the Potts models to provide insights into coupled (paired) mutations, which may arise due to the proximity of the primary and secondary mutations within the three-dimensional protein structure or their participation in a functional role mediated by IN. The output of the data is a plot of double mutant cycles, which provide a biophysical means to interrogate epistasis (*44*) (see **Supplementary Note 1**). We find that the vast majority of mutation pairs (>10^5^) appear close to the center of the distribution (ΔΔE = 0), *i.e*. these pairs are not epistatically interacting. In contrast, most of the pairs of mutations associated with pairwise combinations of E138K + G140A/S + Q148H/K/R are indicated by the Potts model to be amongst the most strongly coupled primary-compensatory mutation pairs observed for IN (**Fig. 2D**). The most strongly coupled pair of IN mutations is G140S/Q148H, followed by G140A/Q148K which is also among the very top pairs. According to the Potts statistical energy analysis, the mutations (S and A) at position 140 interact very strongly with the primary drug resistance mutations (H, K, and R) at position 148 to compensate for the deleterious effects of the primary mutations. Furthermore, the 138K mutation interacts more strongly with the Q148K variant than the Q148H or Q148R variants at position 148, suggesting that E138K plays a stronger role in the E138K/G140A/Q148K pathway than the E138K/G140s/Q148H pathway. In contrast, the mutation pairs at 138 and 140 are not strongly coupled in any of the pathways. This is suggestive that strong couplings connect sites that are likely to co-mutate, allowing for mutations at lesser costs to fitness in patients than the individual mutations alone, and resulting in mutational pathways selected for pathogen escape. Such sets of sites are generally more likely to be associated with resistance to the drugs with which the patients were treated, if resistance cannot be achieved through selectively neutral mutations at single sites, in which case drug treatment would likely be ineffective (*42*).

Collectively, these data can provide insights into the fitness costs associated with single, double, and triple IN mutants, and show which mutations compensate for a loss of fitness caused by primary resistance mutations, and which can potentiate resistance. The Potts statistical analyses, which arise from patient-derived data, provide a means to interpret how pairs of mutations can affect the behavior of the virus in infected individuals undergoing therapy.

### Structural insights into mechanisms of resistance to DTG for the E138K/G140A/Q148K pathway mutants

Measurements done in cells infected with HIV-1 vectors showed that there is a difference in the potency of **DTG** against the IN variants G140A/Q148K and E138K/G140A/Q148K in comparison ot the rest. The relevance of these mutations is underscored by their appearance in cell culture (*27, 45*) and in clinical studies (*23, 46, 47*). We sought to understand the mechanisms underlying these DRMs and the difference in the loss of **DTG** potency using structural biology. We determined high-resolution structures of the HIV-1 intasome constructed from all the possible combinations of single and double mutants that underlie E138K/G140A/Q148K, and the triple mutant, with **DTG** bound. These structures were compared to the structure of the WT HIV-1 intasome with **DTG** bound. We cloned, expressed, and purified all seven mutant INs (E138K, G140A, Q148K, E138K/G140A, E138K/Q148K, G140A/Q148K, and E138K/G140A/Q148K) and assembled intasomes with **DTG** bound. Pooled fractions containing intasomes with **DTG** bound were vitrified on UltraAuFoil gold grids. We collected cryo-EM datasets for all seven mutant assemblies (**table S1**). Each cryo-EM dataset was subjected to iterative 2D and 3D classification and refinement, yielding structures resolved to ~2-3 Å global resolution, with the highest resolution around the active site, where **DTG** is bound (**fig. S1**). The cryo-EM maps demonstrated clear density for all sidechains within the intasome core, the ordered water molecules in and around the active site and, in important cases, the rotameric configurations of the amino acid sidechains. The corresponding atomic models for the complete set of mutants with **DTG** bound was used to develop a detailed understanding the mechanisms of resistance and the contributions of the individual mutations.

The active sites of all experimentally derived mutant structures with **DTG** bound show the substitutions at positions 138, 140, 148, and the drug **DTG** (**Fig. 3A-H**). The structures also show the position of the nearby residue H114, which is not mutated in these structures, but participates in the resistance mechanism (**Fig. 3A**). For all structures, pink arrows point to the mutated residues and asterisks highlight the induced structural changes, including new interactions (red asterisks) or alterations to the structure (yellow asterisks).

**Figure 3:**
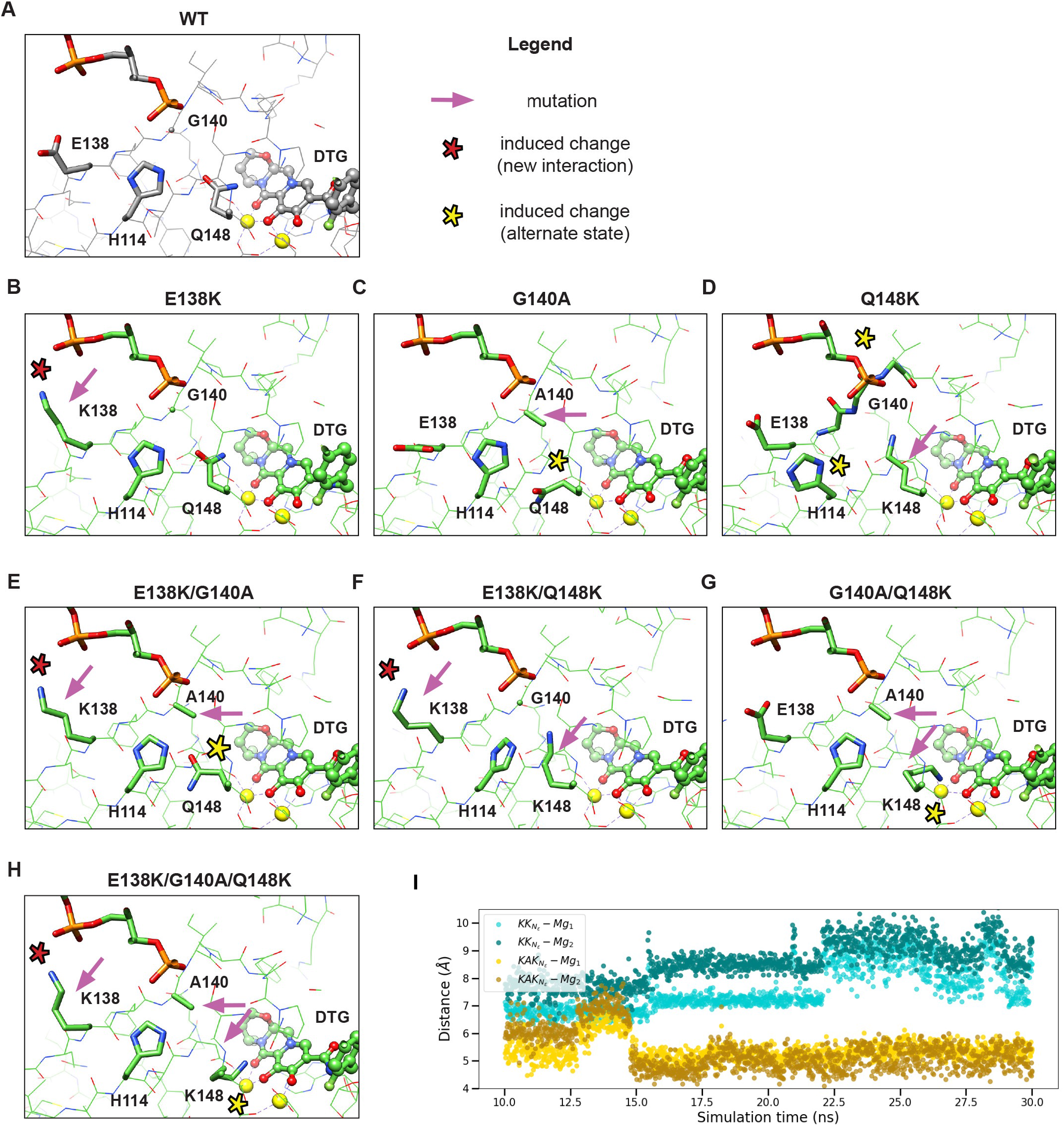
Drug resistant mutations E138K/G140A/Q148K induce altered residue configurations in the active site around bound DTG ligand. Atomic models of the active site from high-resolution intasome (WT and mutant) structures bound to **DTG** are displayed for (**A**) the WT intasome, (**B-D**) the single-mutant variants (**B**) E138K, (**C**) G140A, (**D**) Q148K, (**E-G**) the double-mutant variants (**E**) E138K/G140A, (**F**) E138K/Q148K, (**G**) G140A/Q148K, and (**H**) the triple-mutant variant E138K/G140A/Q148K. In the legend, pink arrows indicate the positions of the mutations, while the asterisks refer to the induced changes, red indicating a novel interaction and yellow indicating an alteration of the configuration of the residue. (**I**) MD simulation of E138K/Q148K (KK) and E138K/G140A/Q148K (KAK) intasomes with **DTG** bound. The atomic distances, measured from the positively charged amino head of Lys148 to either of the Mg^2+^ ions are indicated for the duration of the simulation.

We began by analyzing the structures of the intasomes with single-residue substitutions, E138K, G140A, and Q148K (**Fig. 3B-D**). In the E138K_DTG structure, the ζ-Nitrogen of K138 sidechain approaches within ~3 Å of the backbone phosphate of the 5’-end of the vDNA, participating in a salt bridge with the phosphate (**Fig. 3B**). There are no other additional significant changes in the structure. In the G140A_DTG structure, the introduction of a methyl group affects the sidechain orientation of the nearby residue Q148, inducing a metastable preference for an altered rotameric configuration in which there is a change in the rotation of +120° about the γ-carbon (**Fig. 3C**). The presence of density for the alternative sidechain configuration of Q148 can be clearly seen in the cryo-EM map, which is resolved to ~2 Å in this region (**fig. S2**). If the position of the sidechain of Q148 was not altered, the methyl group of A140 would approach the amide moiety of Q148 to within ~2.5 Å, introducing a weak clash that would destabilize the rotamer that is preferred in the WT intasome bound to **DTG**. The most dramatic changes are observed in the structure of Q148K_DTG, in which the sidechain of K148 impacts multiple nearby residues, leading to: (*i*) an outward shift of the backbone spanning residues 139–141 by up to 1.5 Å, with an RMSD of 1.3 Å between residues 139-141 compared to WT; (*ii*) an induced change in the rotameric configuration of the sidechain of the neighboring residue H114 (**fig. S3** and **Fig. 3D**). Collectively, in the presence of bound drug, the most significant changes in the structure are mediated by Q148K. In contrast, G140A and E138K cause less dramatic changes.

We next analyzed the structures of the double-mutants E138K/G140A, E138K/Q148K, and G140A/Q148K (**Fig. 3E-G**). In the structure of E138K/G140A_DTG, the changes are largely additive to those found in the corresponding single-mutants. The E138K mutation induced salt bridge with the vDNA 5’-phosphate and G140A induced a rotameric reconfiguration of the side chain of Q148; there was density showing an alternative configuration of the sidechain of Q148 (**fig. S2**). In the E138K/Q148K_DTG structure, the E138K mutation again leads to the salt bridge with vDNA, but the other changes induced by Q148K were less prominent, specifically: (*i*) the backbone spanning residues 139-141 largely reverts to the position seen in the WT intasome, with only minor differences in orientation with respect to WT (RMSD 0.3 Å between residues 139-141); (*ii*) the rotameric configuration of H114 is similar to what was seen in the WT, albeit with a minor rotation of the imidazole ring evident in the double mutant. In turn, the sidechain of K148 is oriented slightly toward H114 to avoid clashes (**fig. S3** and **Fig. 3F**). Lastly, in the G140A/Q148K_DTG structure, there are changes that would directly affect the binding of **DTG**. The G140A substitution, as has already been discussed, influences the configuration of the residue at position 148. However, because the residue at position 148 is now Lys, steric effects cause the long sidechain to reorient +240° about the g-carbon, positioning the positively charged head group within 2.7 Å of catalytic Glu152, sufficiently close to form a strong salt bridge interaction, and within 3.6 Å from Mg^2+^ at position A and 3.8 Å from Mg^2+^ at position B (**Fig. 3G** and **fig. S4**). This reorientation, in turn, redistributes the charge around the catalytic metal ions and would, collectively, weaken the chelation by the bound **DTG** (*28*).

Finally, we analyzed the structure E138K/G140A/Q148K_DTG containing all three substitutions (**Fig. 3H**). As with both the single- and double-mutants, E138K makes a salt bridge with vDNA. The remaining substitutions, G140A and Q148K, led to changes in the structure that were similar to what was seen for the G140A/Q148K double mutant, but with a small change in the orientation of the K148 sidechain (**fig. S4**) and an increase in the distance of the Lys to the two Mg^2+^ ions. In the E138K/G140A/Q148K_DTG structure, the positively charged head group is 4.1 Å from Mg^2+^ at position A and 4.4 Å from Mg^2+^ at position B.

The structures suggest that the G140A mutation potentiates resistance through its effect on Q148K. To gain further insight into this interaction and probe the role of G140A in the mechanism of drug resistance, we performed free energy perturbation (FEP) simulations focusing on the effect of the G140A mutation (E138K/Q148K à E138K/G140A/Q148K) on the potency of the INSTI **DTG** using an all-atom molecular dynamics (MD) free energy model. In the simulations, we observe that, when the residue at 140 is Gly (i.e. E138K/Q148K), the positively charged amino head group of Lys148 fluctuates between ~7.6 to ~8.5 Å from the two Mg^2+^ ions. In contrast, when the residue at 140 is Ala (i.e. E138K/G140A/Q148K), there is a corresponding change in the rotamer state of the K148 side chain, and the head group approaches the two Mg^2+^ ions, where its position is stable for the duration of the simulation (**Fig. 3I**). This approach, which is predicted from simulation and without underlying knowledge of the experimental conformation of Q148K in the structure of E138K/G140A/Q148K (see **Supplementary Note 2** and **Methods**), supports our structural biology findings and would, in turn, affect the chelation of the Mg^2+^ ions by **DTG**. Furthermore, based on the FEP calculations, the introduction of the mutation G140A in the background of E138K/Q148K is predicted to reduce the binding affinity of **DTG** by ~1.0 kcal/mol (**fig. S5** and **table S2**), which would translate to a ~10-fold reduction in the EC_50_ of **DTG** when comparing E138K/Q148K with E138K/G140A/Q148K (*48*); this exact reduction in EC_50_ is observed experimentally via virology assays in **Fig. 1B**. In contrast, if the ligand is replaced with **4d**, the change in free energy is within the noise limit (**fig. S5** and **table S2**), which is again consistent with experimentally measured EC_50_ values in **Fig. 1B**. Collectively, the all-atom free energy model simulations successfully account for the effect of the G140A mutation on the change in the binding of **DTG** to intasomes and highlight how G140A mediates the conformation of Q148K, leading to polarization of the Mg^2+^ ions and weakening chelation by the bound ligand.

Taken together, the structures and the simulations provide a framework for understanding, at an atomic level, the changes that are observed in the mutations within the E138K/G140A/Q148K pathway when **DTG** is bound. The E138K mutation induces a salt bridge with the 5’-end of the vDNA, an interaction that is observed in all structures. This interaction stabilizes the position of the 5’-end of the vDNA, and presumably the conformation of the active site, which explains its mild compensatory behavior observed in enzymology and virology assays (**Figs. 1–2**). The Q148K mutation can adapt distinct configurations in the active site because the Lys sidechain is flexible and dynamic, and the exact configuration of the Lys sidechain is strongly affected by the neighboring residues. By itself, Q148K points toward the backbone residues 139-141 and induces multiple, presumably deleterious, changes to the local structure. The accessory mutations E138K or G140A largely (but not completely) restore the WT configuration of the protein backbone at residues 139-141 and the rotameric configuration of the sidechain of H114. However, they do so in distinct ways: the mutation E138K, through its interaction with vDNA, pulls the peptide “back into position”, leading to the reconfiguration of both the backbone and the H114 sidechain. In contrast, A140 relieves steric stress by pushing K148 toward the two Mg^2+^ ions. Notably, it is the G140A mutation that most affects the orientation of the sidechain at position 148, and thus the mechanism of drug resistance. If residue 148 is Lys, the mutation G140A leads to a repositioning of the sidechain by +240° about the γ-carbon, bringing the ζ-Nitrogen to ~3.6–4.4 Å of the two Mg^2+^ ions. This reconfiguration, in turn, polarizes the charge distribution around the metal ions and weakens the chelation with the bound **DTG**, reducing its binding, and consequently, leading to drug resistance.

### Mechanism of resistance of the Q148H/K/R mutants

We next asked how each of the three different amino acid mutations Q148H/K/R affect the binding of INSTIs. In prior work, the SIVrcm IN was used as a model for HIV-1 IN, and the analysis was limited to the double mutant G140S/Q148H with **BIC** bound (*28*). Here, we determined structures of the HIV-1 intasomes E138K/G140A/Q148R and E138K/G140S/Q148H, with **DTG** bound, resolved to 2.5 Å and 2.2 Å, respectively (**table S1**). These structures were compared to the structure of E138K/G140A/Q148K intasome with **DTG** bound and to the structure of the G140S/Q148H SIVrcm intasome with **BIC** bound.

The most notable observation in these structures is the relative position of the sidechain at position 148 and its relationship to the two Mg^2+^ ions. In the structure of the E138K/G140A/Q148K intasome with **DTG** bound, K148 approaches the Mg^2+^ ions to within ~4.1– 4.4 Å, as discussed above (**Fig. 4A**). In the structure of E138K/G140A/Q148R intasome with **DTG** bound, the guanidium group resides in a similar overall configuration relative to the Mg^2+^ ions, but the sidechain is approximately ~2.5 Å longer, which positions the head group underneath the bound INSTI and further from the Mg^2+^ ions (5.2 Å), as measured from the ζ-carbon where the positive charge is localized (**Fig. 4B**). The longer sidechain affects the position of residues 144-147, leading to a small local displacement of this segment (the RMSD is 0.3 Å relative to WT), **fig. S6**). The displacement of residues 144-147 is similar when the E138K/G140A/Q148R intasome is compared to both E138K/G140A/Q148K (RMSD 0.3 Å) and E138K/G140S/Q148H (RMSD 0.4 Å). The Q148R mutation causes a weakening of the chelation by **DTG**, which, when combined with other mutations, also weakens the binding of the drug. In contrast to the two structures described above, the sidechain of His is shorter in the E138K/G140S/Q148H intasome and, accordingly, the His sidechain is also furthest from the Mg^2+^ ions, (**Fig. 4C**).

**Figure 4:**
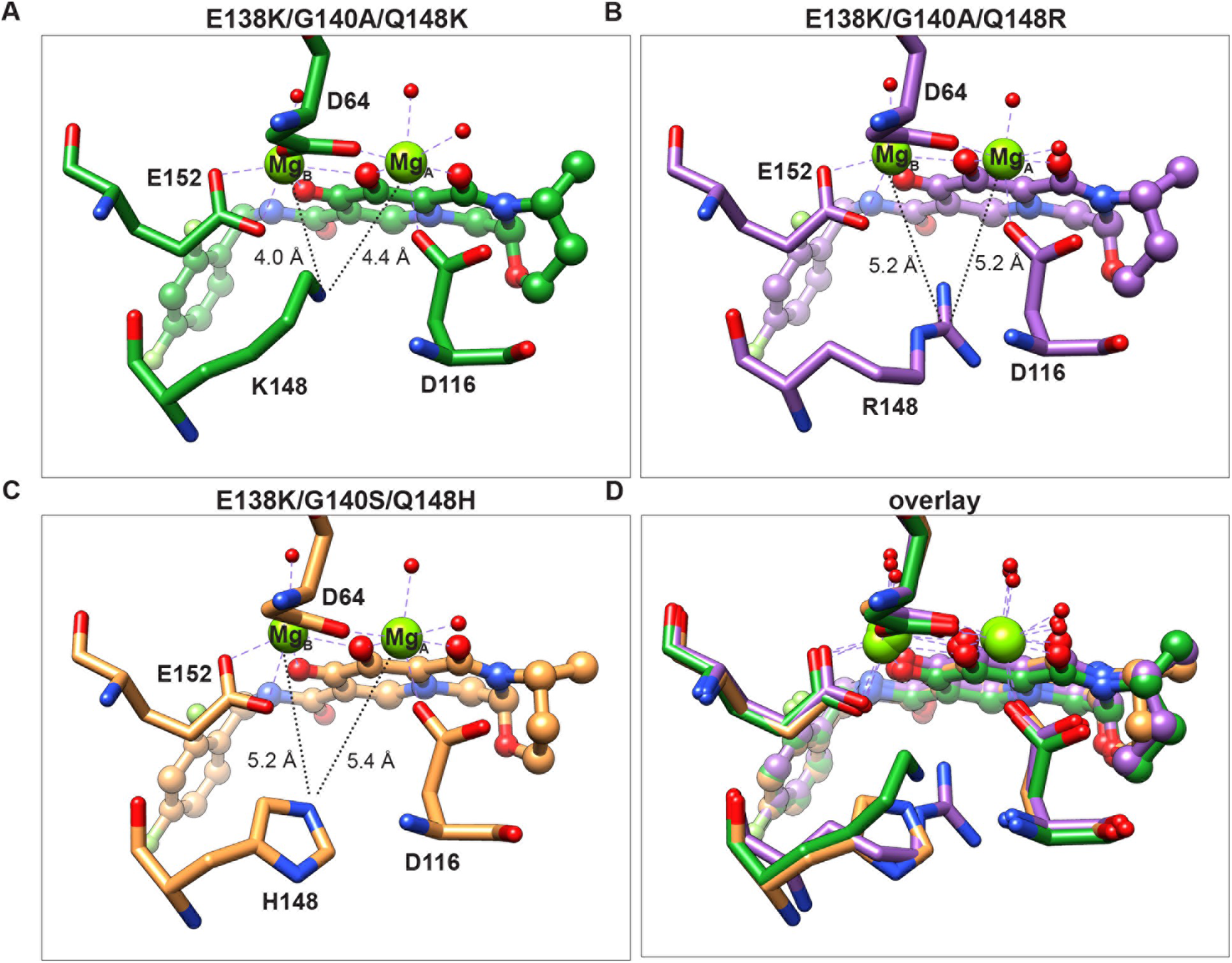
Structural insights from comparisons of Lys, Arg, and His mutations at position 148. Atomic models of the region surrounding the bound **DTG** from experimental structures of (**A**) E138K/G140A/Q148K, (**B**) E138K/G140A/Q148R, and (**C**) E138K/G140S/Q148H intasomes with **DTG** bound. (**D**) Overlay of the three experimentally determined structures. Relevant atomic distances are indicated in the snapshots.

To compare the distinct model systems used to examing INSTIs, we next compared the structure of the E138K/G140S/Q148H HIV-1 intasome with **DTG** bound to the structure of G140S/Q148H SIVrcm intasome with **BIC** bound, which was solved previously (*28*). There are small but significant differences in the two structures. The RMSD of the backbone is 0.5 Å between residues 136-150 and 0.6 Å between residues 110-116, which surround the mutated residues (**fig. S7A-C**). In addition, there are nine residues within 10 Å of the bound drugs that differ between HIV-1 and SIVrcm (**fig. S7D**). If the E138K mutation is present in the HIV-1 intasome, the sidechain of H114 rotates toward the S140/H148 pair, inducing an alternate rotamer in S140 compared the SIVrcm intasome. This causes the Nε2 atom of H114 to approach closer to the Nδ1 atom of H148 by 1.0 Å in the HIV-1 intasome. The repositioning of H114 strongly implicates this residue in the resistance mechanism through its interaction with H148, which would facilitate proton transfer to stabilize the buildup of positive charge around the Mg^2+^ ions. The relative positions of the side chains of the mutant residues and their proximity to the Mg^2+^ ions can be seen in an overlay of the three structures (**Fig. 4D**).

Collectively, these data provide mechanistic insights into how the distinct residue changes Q148H/K/R affect **DTG** binding. The proton donating capabilities of the three sidechains are Arg à Lys à His. Whereas Arg is the most potent proton donor, it is also more distant from the Mg^2+^ ions in comparison to Lys, which explains why its effect on resistance, at least in the context of accessory mutations E138K/G140A, is less than that of Lys at the same position. The His-mediated polarization of Mg^2+^ ions is expected to be weakest, and its effect on the binding of INSTIs is also expected to be less. Thus, it is both the position and the charge of the sidechains at residue 148 relative to the Mg^2+^ ions that determines the effects of the mutation on drug binding. Given that the chemical scaffolds of **DTG** and **BIC** are similar, as are the inhibitory potencies, our findings can reasonably be extended to **BIC**. We also expect similar rules to apply to **CAB**, although the effects will be more pronounced, because this drug is less potent against many DRMs (*22*). Furthermore, as changes to residues G140A/S+Q148H/K/R are some of the most prevalent in INSTI-resistant HIV-1 IN (*23*), the current structures should also provide a valuable foundation for future drug design efforts aimed at combatting resistance.

### Stacking of the terminal adenine base of vDNA on the INSTI contributes to the potency of compound 4d against the E138K/G140A/Q148K family of DRMs

We next wished to determine why the naphthyridine-containing compound **4d** maintains potency against most of the DRMs we tested, in particular the triple mutant E138K/G140A/Q148K, when **DTG** does not. We thus determined the structure of the E138K/G140A/Q148K mutant intasome from HIV-1 with bound **4d** at 2.75 Å resolution and compared it with the structure of the same mutant with **DTG** bound (**table S1**).

We first compared the configuration of the ligand when either **DTG** or **4d** is bound to E138K/G140A/Q148K HIV-1 intasomes. Both compounds contain a di-fluorinated benzene that makes a π-π stacking interface with the base of the penultimate cytosine of the vDNA. The fluorinated benzene is connected to the INSTI core through an amide-containing linker that is identical in both **DTG** and **4d**, and contacts between the first ring of both ligands to the intasome are similar. The major differences between the compounds are in the way they chelate Mg^2+^ ions and the nature of the extension at the solvent-end of the active site (**Fig 5A-C**). Whereas **DTG** uses three oxygen atoms – all external to the ligand core – to coordinate the two Mg^2+^ ions (**Fig. 5A**), **4d** has a nitrogen embedded into its naphthyridine core (**Fig. 5B**). This leads to an overall ~10° rotation of **4d** toward the Mg^2+^ ions (**Fig. 5C**). **4d** has a hexanol moiety appended to the 6-position of the naphthyridine scaffold instead of the 4-methyl-1,3-oxazinane ring of **DTG**. This hexanol displaces weakly bound waters in the apo structure and extends out toward the solvent-exposed cleft, sandwiched between N117-G118 and P142-Y143, a configuration that is expected to stabilize the binding of the ligand to HIV-1 intasomes containing DRMs. Furthermore, when **4d** is bound to E138K/G140A/Q148K intasomes, relative to the structure with **DTG** bound, Tyr143 extends outward by 1.5 Å, a result of the hexanol extension making Van Der Waals contacts in this region (**Fig. 5C**).

**Figure 5:**
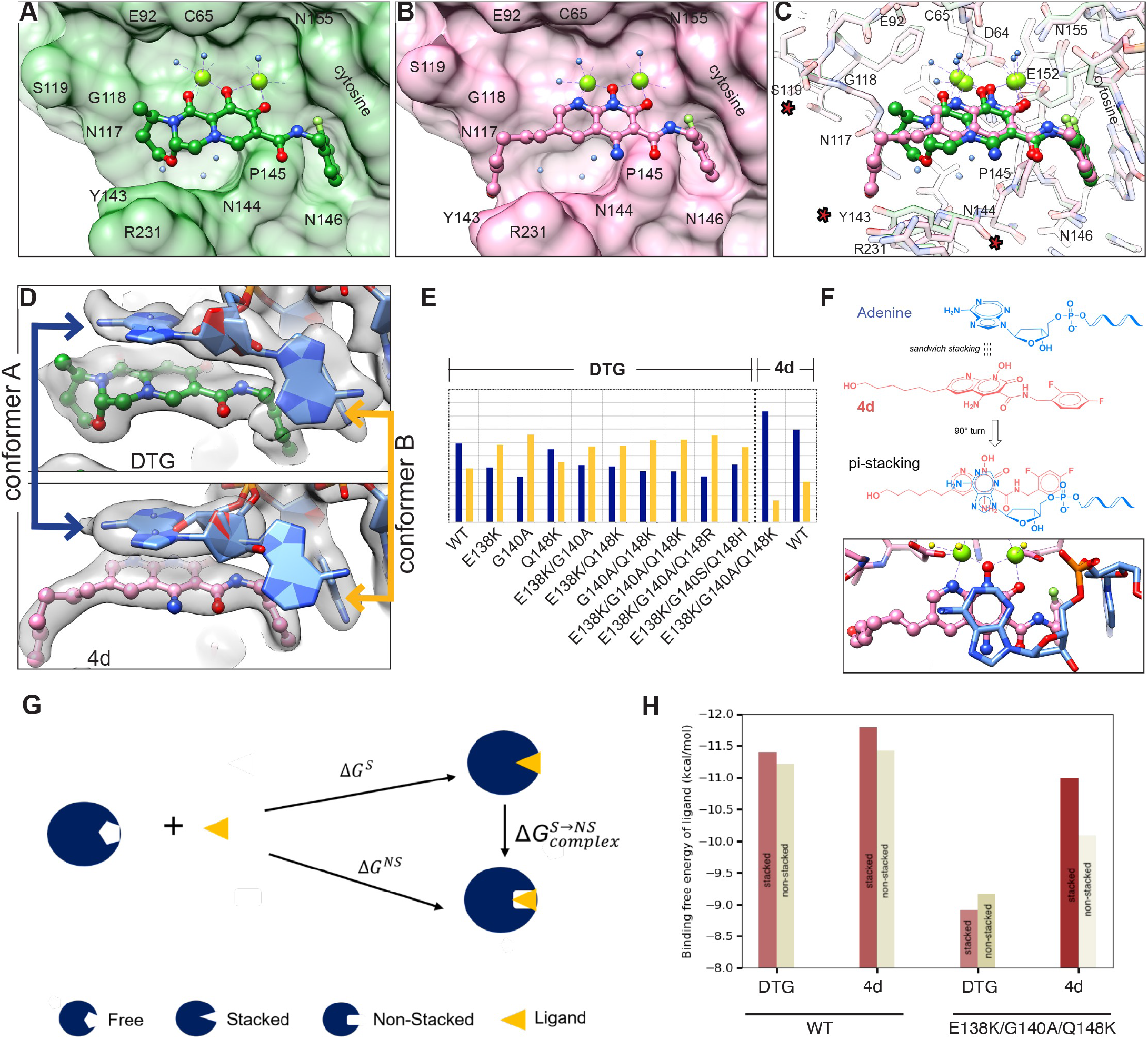
Structure-based explanation for 4d potency. (**A-B**) Atomic models of (**A**) **DTG** or (**B**) **4d** from high-resolution structures of the E138K/G140A/Q148K intasome bound to either of the two drugs, respectively. The intasome is displayed as a surface view, while the drugs are displayed in ball-and-stick. The terminal adenine base is removed, for clarity. (**C**) Overlay of the two structures in A-B, with all atoms displayed as ball-and-stick. Residues whose positions vary between the two structures are highlighted with a red asterisk. (**D**) Close-up of the cryo-EM density of the terminal adenosine and its interaction with the bound drug. When the intasome contains the E138K/G140A/Q148K mutation, experimental density is weak for conformer A, which stacks on top of the bound INSTI, when the ligand is **DTG**, but is pronounced when the ligand is **4d**. (**E**) Quantitative comparison of the rotameric occupancies of the two conformers, A and B, in all structures derived in the current work. Only the structures in which **4d** is bound clearly show pronounced conformer A occupancy. (**F**) Chemical schematic of the π-π stacking between the terminal adenine base and **4d**, which helps to maintain the bound drug in the active site of the intasome. (**G**) Thermodynamic cycle used for estimating the ligand binding free energies to the target in the two conformers. (**H**) Estimates of the free energies for the stacked (brown) and the non-stacked (tan) configurations of the terminal adenine base, for WT (left) and E138K/G140A/Q148K (right) DRM intasomes.

We next examined the structural configurations within the intasome with either **DTG** or **4d** bound. There are no significant differences in the position of the mutated amino acids E138K, G140A, or Q148K, and the free amine on the sidechain of K148 occupies a similar position (<0.2 Å difference) relative to the Mg^2+^ ions (**fig. S8A**). There are minor variations in the two models in the positions of S119, R228, R231, and other small amino acids, but the significance of these ancillary changes is unclear (**Fig. 5C**). There is some rearrangement of the hydration shell around the ligands, while the three core bound waters underneath the ligand, and the fourth water, which is displaced if the mutation Q148K is present, are entirely conserved (**fig. S8B**). The major differences, at the level of the intasome, are in the configuration of the terminal adenosine of the vDNA. This adenosine, which is universally conserved among retroviral INs, can adapt two distinct conformations in the active site: (*i*) one that stacks the adenine base on the bound INSTI, and (*ii*) one in which the base extends freely into the space between the IN subunits. Quantification of the relative occupancies of each conformation revealed a strong preference for the stacked conformation when **4d** is bound (**Fig. 5D**). A similar stacked conformation is a minority population in all mutant structures with **DTG** bound (**Fig. 5E**), suggesting that the high fractional occupancy of the stacked conformation when **4d** is bound is specific to this INSTI and helps maintain the drug bound in its preferred position in the presence of the DRMs.

Stacking of the terminal adenine establishes a π-π bond reminiscent of the one between the halogenated benzene and the base of the penultimate cytosine (Fig. 5F). The stronger stacking of **4d** with the terminal adenosine is expected to stabilize the binding of **4d** relative to **DTG**. We thus wished to estimate how stacking affects the potency of **4d**. To this end, we can use a simple thermodynamic cycle as shown in Fig. 5G. In this schematic, the ligand is either **DTG** or **4d**, and the target is the adenine base; the ligand binding free energies for the stacked configuration ΔG^S^, the non-stacked configuration ΔG^NS^, and the transition ΔG^S→NS^, complete a thermodynamic cycle that describes the relationships between the distinct conformational states. In the context of this thermodynamic cycle, the stacking probabilities of **4d** and **DTG**, determined from the cryo-EM maps, and the EC_50_ values, determined via virology assays, provide experimental constraints that can be used to solve a system of equations to yield binding affinities in the stacked and in the non-stacked conformations for **4d** and **DTG** to the E138K/G140A/Q148K triple mutant (see Supplementary Note 3). The inputs, and the resulting binding affinities of **4d** and **DTG** to the stacked and non-stacked forms of the E138K/G140A/Q148K intasome complex are listed in table S3. We find that, while the binding free energies of the adenine-**DTG** conformers to the E138K/G140A/Q148K intasome are similar (both stacked and non-stacked are ~ −9 kcal/mol), the stacked adenine-**4d** conformer shows a more favorable binding free energy of ~ −11 kcal/mol, and this conformer makes the dominant contribution to the binding strength (Fig. 5H). Furthermore, the difference in binding free energy of ligand between **4d** and **DTG** for the E138K/G140A/Q148K intasome complex is higher in the stacked conformation (~ −2.1 kcal/mol) versus the non-stacked conformation (~ −0.9 kcal/mol), indicating that stacking is especially relevant in the context of the E138K/G140A/Q148K DRM (table S3). This observation is consistent with structural biology, wherein the conformation of **4d**, in the context of the E138K/G140A/Q148K intasome, is displaced toward the adenine base, which is caused by the Q148K mutation “underneath” the ligand; this same displacement is not observed when **DTG** is bound (Fig. S9). Thus, adenine stacking onto **4d** is a strong contributing factor to the increased potency of **4d** as compared with **DTG** and should be considered in future design strategies to optimize INSTI binding to DRM intasomes.

## DISCUSSION

### Mechanistic insights into loss of DTG potency revealed through mutations containing G140A/Q148K

Our efforts to probe mutations E138K + G140A/S + Q148H/K/R revealed specific weaknesses in the ability of **DTG** to inhibit certain complex DRMs. Specifically, if the G140A mutation is present in combination with Q148K, as in the double mutant G140A/Q148K or the triple mutant E138K/G140A/Q148K, the inhibitory potency of **DTG** drops by ≳1 order of magnitude (**Fig. 1**). Thus, we wished to define the mechanism by which viruses containing DRMs G140A + Q148K become resistant to **DTG**. Our structures explain why high-level resistance to **DTG** requires the G140A mutation, and its specific pairing with Q148K. Loss of **DTG** potency can be primarily attributed to the introduction of positive charge around the Mg^2+^ ligands by K148. Although E138K can potentiate resistance, its effects are modest. In contrast, G140A strongly potentiates resistance. This reduction in potency stems from the effect that G140A has on inducing a reorientation of K148 toward the Mg^2+^ ions and charge redistribution in the surrounding region (**Fig. 3**). Thus, the proximity of the charged ζ-amine on K148 to Mg^2+^ is directly related to the effect of A140. Based on inhibitory potencies, the same logic can be extended to the mutation G140S.

The structural changes induced by K148 potentiate resistance, but they also incur a fitness cost. For a resistant virus to be selected by a drug, the gain from the development of resistance must be greater than the loss incurred by the reduction in replicative capacity. Both the infectivity of the virus and enzymatic activity of IN are substantially reduced in viruses containing the Q148K mutation. This is readily explained; the bulky charged Lys at position 148 intrudes into the active site of IN, impairing its activity (**Fig. 2A-C**). Despite their detrimental effect on fitness, DRMs containing G140A/Q148K combinations appear in patients (*23, 46, 47*) demonstrating their clinical relevance. Taken together, the changes induced in the structure of the intasome make viruses containing G140A/Q148K or E138K/G140A/Q148K mutations less susceptible to inhibition by **DTG**, but still being able to mediate integration of vDNA into host DNA, explaining why they are found in both cell culture and PLWH.

### The proximity of the amino acid sidechain to the Mg^2+^ ions and the strength of the proton donating capabilities of the head group defines the potency of resistance for Q148H/K/R variants

There are three possible mutations at position 148: His, Lys, or Arg. Variants containing Lys, including the double mutant G140A/Q148K and the complex mutant E138K/G140A/Q148K, are the least susceptible to inhibition by **DTG** (**Fig. 1**). Structural biology shows that the drop in susceptibility to **DTG** can be primarily explained by the proximity of the NH2 moiety of the flexible Lys sidechain to the two Mg^2+^ ions. In the presence of bound **DTG**, Lys is positioned closer than both the Arg and the His variants by ~1.0 Å (**Fig. 4**), which explains the ~10-fold loss of **DTG** potency in the presence of G140A/Q148K mutations. Although Arg is further from the Mg^2+^ ions, it is a stronger proton donor, and thus the Arg would also be expected to have a significant effect on the polarization around the Mg^2+^ ions, weakening the chelation by the bound drug. Indeed, we observe loss of potency of **DTG** against the complex mutation E138K/G140A/Q148R (**Fig. 1**). We do not observe loss of potency of **DTG** against G140A/Q148R intasomes, which implies that the Arg is slightly reoriented in the double mutant in comparison to the triple mutant. If the INSTIs under consideration are **DTG** and **4d**, the least effective substitution at position 148 is His. Both compounds potently inhibit all combinations of DRMs (up to three changes) containing the Q148H mutation (**Fig. 1**). This is consistent with the larger distance from the sidechain of His to the Mg^2+^ ions. In sum, it is the specific configuration of the sidechain of the residues relative at position 148 to the two Mg^2+^ ions that determines the effects of the mutation on ligand binding.

Despite their relatively mild effect on ligand binding, variants containing Q148H arise and are prevalent in patients being administered either first or second-generation INSTIs (*23, 46, 47*). The broad persistence of Q148H variants may be, at least in part, due to the fact that they are readily selected by the first generation INSTIs, which have been used clinically since 2007. The propensity to select mutants that carry the G140S/Q148H mutations may be due to the relatively high replicative capacity and enzymatic activity of the G140S/Q148H variants in the absence of any drugs, supported here by *in vitro* enzymatic and single cycle replication assays (**Fig. 2A-C**). Thus, while the strength of the positive charge and the proximity to the Mg^2+^ ions can explain the potency of resistance for the Q148H/K/R group variants, their appearance in the clinical setting and persistence in patient populations depends on which INSTIs are used to select the mutants, on the accompanying mutations at positions 138 and 140 (among other positions), and on how the different mutants affect the complex interplay of these two factors with the collection of viruses present in the infected individual. Given that Q148K/R mutants are less susceptible to inhibition by **DTG**, we would expect mutants containing Q148K/R to begin appearing more frequently in patients being administered **DTG**, and by extension, other 2^nd^ generation INSTIs.

### Preferences for G140A/S are influenced by the mutational context of Q148H/K/R

The residue variant at position 140, either Ser or Ala, plays an important role in modulating resistance. Based on “intrinsic” fitness constraints identified using Potts double-mutant-cycle values, which reflect internal constraints in the enzyme that apply in any environment in the Potts framework, the most strongly interacting pair of IN mutations is G140S/Q148H, which is followed by G140A/Q148K, and then the cluster of residue pairs that include G140S/Q148R, G140S/Q148K, G140A/Q148R, and G140A/Q148H (**Fig. 2D**). Our data, generated using multiple experimental systems to measure inhibitory potencies, fitness, enzyme activity, and atomic structure, helps to explain why S140 appears to arise in conjunction with Q148H, whereas both A140 and S140 can arise in conjunction with Q148K/R. In the case of the double mutant G140S/Q148H, based on structure, S140 directly interacts with H148 via the polar hydroxyl group of Ser and the Nδ1 atom of His. As previously suggested (*28*), this interaction may potentiate resistance by contributing to the proton donating capability of this pair of histidines, although our data suggests that this effect is weak (**Fig. 1**). More importantly, the presence of G140S restores enzymatic activity and viral fitness (**Fig. 2A-C**), supporting its previously described role as a compensatory mutation (*33*). The compensatory effect appears to be caused by S140 forming hydrogen bonds with both H148 and T115, which creates a favorable energetic environment around the active site when the mutation Q148H is present in conjunction with G140S. In contrast to G140S, the mutation G140A does not have free protons to donate and it should not contribute favorably to the energetic environment. Consistently, G140A, by itself or in combination with other DRMs, generally does not restore enzymatic activity or viral infectivity (**Fig. 2A-C**). Instead, it modulates the conformation of residue 148 when a drug is bound, positioning the residue closer to the Mg^2+^ ions. This is seen in the structures of Q148K-containing intasomes (**Fig. 3**) and, based on comparisons of inhibitory potencies, is implied for Q148R-containing intasomes. Because the mutation pairs G140S/Q148K, G140A/Q148R, and G140S/Q148R arise nearly as frequently as the mutation pair G148A/Q148K (**Fig. 2D**), and there are no distinct groupings that can be discerned based on measurements of inhibitory potencies, enzyme activity, or viral replicative capacities, we conclude that, in the presence mutations Q148K/R, the variant G140S should function analogously to the variant G140A. Collectively, these data indicate that the selection of G140A/S depends on the substituent at position 148 and that preferences for certain combinations of mutations can, to some extent, be explained by the structures. However, a complete understanding of exactly how the various combinations of mutations interact and how such interactions will affect the structure and function of IN would require considering the complex interplay between the virus inhibition (and the specific INSTIs used to treat PLWH), the background mutational landscape of the virus, and the relative fitness of the viruses in the presence of the different drugs.

### Mechanism of action of E138K in drug resistance and functional compensation

The E138K mutation can reduce the potency of **DTG** and, in at least some mutant backgrounds, plays a compensatory role, restoring some enzyme activity and viral fitness. Prior work suggested that the similar mutation E138T in SIVrcm potentiates resistance by serving as an H-bond donor to His114 within an extended network that reinforces the positive charge around **DTG**, weakening its chelation of Mg^2+^ (*28*). However, the role of the E138K mutation in potentiating resistance appears to be different, based on multiple observations. First, the distance between Lys138 and His114 always ranges from ~6—8 Å, which is too large to form an H-bond network. Second, the mutation G140A, which can arise in conjunction with E138K, lacks an H-bond acceptor, but the aliphatic sidechain of Ala, like Ser, strongly reduces the potency of **DTG** (**Fig. 1**). Both observations would preclude the establishment of an H-bond network. Instead, we suggest that the mutation E138K leads to micro-rearrangements of the protein and solvent which, collectively, lead to small but significant changes to the active site. The compensatory behavior of E138K may stem from stabilizing the binding and/or proper positioning of the vDNA. The strand transfer step of integration is a dynamic process that requires careful coordination of multiple substrates and the sequestration of the 3’-OH moiety on the vDNA for SN2-mediated nucleophilic attack (*5, 49*). Many mutations are known to affect the strand transfer reaction and, presumably, the conformation and dynamics of the vDNA. A salt bridge between Lys138 and the vDNA phosphate backbone should help stabilize the position of the 5’-end of vDNA, especially when other DRMs are present. Our data shows that the E138K mutation could be selected in the presence of several different mutant backgrounds, suggesting that this is a general compensatory mechanism that may be encountered when multiple DRMs accumulate.

### Potency of 4d and the naphthyridine compounds

Loss of potency of INSTIs that is caused by mutations at positions E138K+G140A/S+Q148H/K/R can be addressed through the development of better compounds. Select chemical modifications on the ligand, such as the primary amine at the 4-position of the naphthyridine scaffold (*50*), allow **4d** to retain better binding and potency than **DTG** (*21*). Such prior observations suggested that introducing specific chemical groups could enhance interactions with the intasome to minimize resistance mediated by the Lys or Arg variants at position 148. In this regard, the consistent observation of stacking mediated by the terminal adenine base of the vDNA, and its predominance when the INSTI **4d** is bound was encouraging. The fact that the stacking is more pronounced when **4d** is bound than when **DTG** is bound (and by extension, also **BIC** and **CAB**) is chemically consistent with the fact that the π-π interactions are expected to be stronger against the naphthyridine moiety of **4d** than against the tricyclic moiety of **DTG**. Although stacking is not the only contributor to potency, it appears to be an important one. Likely, there are additional means by which to exploit, and perhaps enhance, the stabilization of the terminal adenine base, to generate even tighter binders. It is also relevant to identify novel means to counteract loss of INSTI potency against the prevalent group of Q148H/K/R mutations that affect Mg^2+^ chelation. Future efforts should explore these ideas and aim to design novel compounds that are broadly effective against drug resistant HIV.

## Author contributions

D.P., D.L., M.L., and R.C. conceived the work and designed the initial experimental strategies. M.L. developed the conditions for intasome production and purification of the mutant samples. Z.L. contributed to protein purification. D.O.P., M.L., and Z.S., performed cryo-EM data acquisition, image processing, and structure determination. T.S. and D.L. built and refined the models. S.J.S. performed virology assays and helped with the interpretation of the data. M.L. performed enzymatic assays. A.B. and A.H. performed the Potts analyses. Q.S. performed MD simulations. X.Z. synthesized compound **4d** for structural work. D.L. wrote the original draft of the manuscript, and all authors contributed to writing and editing. D.L. provided funding and supervised the work.

## Data and materials availability

The cryo-EM maps and atomic models have been deposited into the Electron Microscopy Data Bank and Protein Data Bank under the accession codes listed in **Table S1**. All data needed to evaluate the conclusions in the paper are present in the paper and/or the Supplementary Materials.

## MATERIALS AND METHODS

### Vector constructs

To make the IN mutants, the vector pNLNgoMIVR-ΔENV.LUC, which has been described previously (*22*), was used to produce the new IN mutants analyzed in this study. The IN open reading frame was removed from pNLNgoMIVR-ΔENV.LUC by digestion with KpnI and SalI and resulting fragment was inserted between the KpnI and SalI sites of pBluescript KS+. Using that construct as the wild-type template, the following HIV-1 IN mutants were prepared using the QuikChange II XL site directed mutagenesis kit (Agilent Technologies, Santa Clara, CA): E138K, G140A, G140S, Q148H, Q148K, Q148R, E138K/G140A, E138K/G140S, E138K/Q148H, E138K/Q148K, E138K/Q148R, G140A/Q148H, G140A/Q148K, G140A/Q148R, G140S/Q148H, G140S/Q148K, G140S/Q148R, E138K/G140A/Q148H, E138K/G140A/Q148K, E138K/G140A/Q148R, E138K/G140S/Q148H, E138K/G140S/Q148K, and E138K/G140S/Q148R. The following sense oligonucleotides were used with matching cognate antisense oligonucleotides (Integrated DNA Technologies, Coralville, IA) in the mutagenesis:

E138K, 5’-TGGGCGGGGATCAAGCAGAAATTTGGCATTCCCTACAAT −3’; G140A, 5’-
GGGATCAAGCAGGAATTTGCTATTCCCTACAATCCCCAA −3’; G140S, 5’-
GGGATCAAGCAGGAATTTTCCATTCCCTACAATCCCCAA −3’; Q148H, 5’-
CCCTACAATCCCCAAAGTCATGGAGTAATAGAATCTATG −3’; Q148K, 5’-
CCCTACAATCCCCAAAGTAAAGGAGTAATAGAATCTATG −3’; Q148R, 5’-
CCCTACAATCCCCAAAGTCGAGGAGTAATAGAATCTATG −3’; E138K for E138K/G140A, 5’-
GGGATCAAGCAGAAATTTGCCATTCCCTACAATCCCCAA −3’; and E138K for E138K/G140S,
5’-GGGATCAAGCAGAAATTTAGCATTCCCTACAATCCCCAA – 3’.

The IN mutants E138K, G140A, G140S, Q148H, Q148K, and Q148R were made using the WT template and the respective oligonucleotide listed above. Because of the complexity of the IN mutants (some IN mutants may have 2 or more IN mutations), certain IN mutants in the KS construct will be used as templates for additional rounds of mutations. The IN mutants E138K/G140A, E138K/G140S, E138K/Q148H, E138K/Q148K, E138K/Q148R, G140A/Q148H, G140A/Q148K, G140A/Q148R, G140S/Q148H, G140S/Q148K, G140S/Q148R, E138K/G140A/Q148H, E138K/G140A/Q148K, E138K/G140A/Q148R, E138K/G140S/Q148H, E138K/G140S/Q148K, and E138K/G140S/Q148R were made as follows. The IN mutant E138K/G140A was made using the previously constructed IN mutant G140A as the template, while the proper E138K oligonucleotide listed above was used to make the second amino acid substitution. The IN mutant E138K/G140S was made using the previously constructed IN mutant G140S as the template, while the proper E138K oligonucleotide aforementioned was used to make the second amino acid substitution. To make the IN mutants E138K/Q148H, E138K/Q148K, E138K/Q148R, the previously constructed IN mutant E138K was used as the template and correct oligonucleotide for either Q148H, Q148K, or Q148R was used to make the second amino acid substitution. To make the IN mutants G140A/Q148H, G140A/Q148K, G140A/Q148R, the previously made IN mutant G140A was used as a template, and the proper oligonucleotide for either Q148H, Q148K, or Q148R was used to make the second amino acid substitution. For the IN mutants G140S/Q148H, G140S/Q148K, G140S/Q148R, the previously constructed IN mutant G140S was used as a template and the correct oligonucleotide for either Q148H, Q148K, or Q148R was used to make the second mutation. For the IN mutants E138K/G140A/Q148H, E138K/G140A/Q148K, E138K/G140A/Q148R, the previously constructed IN double mutant E138K/G140A was used as a template and the proper oligonucleotide for either Q148H, Q148K, or Q148R was used to make the third amino acid substitution. For the IN mutants E138K/G140S/Q148H, E138K/G140S/Q148K, and E138K/G140S/Q148R, the previously made IN double mutant E138K/G140S was used as the template and either the proper oligonucleotide for Q148H, Q148K, or Q148R was used to make the third mutation.

After the mutagenesis, the DNA sequence of each construct was determined. The mutated IN coding sequences from pBluescript KS+ will then be subcloned into pNLNgoMIVR-ΔEnv.LUC (between the KpnI and SalI sites) to produce mutant HIV-1 constructs, which were also checked by DNA sequencing.

### Cell-based replication assays

Cell-Based assays. HIV-based viral vectors with either WT or mutant IN were used in single-round infectivity assays to determine the antiviral activity (EC_50_ values) of the compounds as previously described (*51*). Briefly, VSV-g-pseudotyped HIV-1 was produced by transfections of 293 cells with pNLNgoMIVR^-^ΔLUC and pHCMV-g (obtained from Dr. Jane Burns, University of California, San Diego) using the calcium phosphate method. At approximately 6 hrs after the calcium phosphate precipitate was added, 293 cells were washed twice with phosphate-buffered saline (PBS) and incubated with fresh media for 48 hrs. The virus-containing supernatants were then harvested, clarified by low-speed centrifugation, filtrated, and diluted for preparation in antiviral infection assays. On the day prior to the screen, HOS cells were seeded in a 96-well luminescence cell culture plate at a density of 4000 cells in 100 *μ*L per well. On the day of the screen for antiviral activity infection assays, cells were treated with compounds from a concentration range of 5 *μ*M to 0.0001 *μ*M using 11 serial dilutions and then incubated at 37°C for 3 hrs. After compound incorporation, 100 *μ*L of virus-stock diluted to achieve a luciferase signal between 0.2 and 2.0 Relative Luciferase Units (RLUs) was added to each well and further incubated at 37°C for 48 hrs. Infectivity was measured by using the Steady-lite plus luminescence reporter gene assay system (PerkinElmer, Waltham, MA). Luciferase activity was measured by adding 100 *μ*L of Steady-lite plus buffer (PerkinElmer) to the cells, incubating at room temperature for 20 mins, and measuring luminescence using a microplate reader. Antiviral activities were normalized to the infectivity in cells that featured the absence of target compounds. KaleidaGraph (Synergy Software, Reading, PA) was used to perform non-linear regression analysis on the data. EC_50_ values were determined from the fit model.

### Single-round infection assays

A modified version of the single-round infectivity assay was used to determine the replication of the INSTI-resistant mutant vectors. Briefly, 200 ng of a WT or INSTI-resistant mutant HIV-1 based vector was added to each well in 96-well plates, incubated for 48 hrs, and luciferase activity was measured as mentioned above. The luciferase activity of the WT virus was set to 100%, and the infectivity of the mutant viruses was measured as a percentage of WT (*20*).

### IN protein expression and purification

His-tagged Sso7d-IN and mutants were expressed and purified essentially as described before (*14, 15, 34*). Briefly, the integrase was expressed in *E. coli* BL21 (DE3) and the cells were lysed in buffer containing 20 mM Hepes pH 7.5, 10% glycerol, 2 mM 2-mercaptoethanol, 20 mM imidazole and 1M NaCl. The protein was purified by nickel-affinity chromatography and the His-tag was removed with thrombin. Aggregated protein was removed by gel filtration on a HiLoad 26/60 Superdex-200 column (Cytiva) equilibrated with 20 mM Hepes pH 7.5, 10% glycerol, 2 mM DTT, 1 mM EDTA and 0.5 M NaCl. The protein was concentrated using an Amicon centrifugal concentrator (EMD Millipore) as necessary, flash-frozen in liquid nitrogen and stored at −80°C. His-tagged LEDGF-IBD (307-460) was expressed and purified in the similar way except His-tag was not removed from IBD.

### Intasome assembly and purification

The double-stranded vDNA substrate was prepared by annealing the following oligonucleotides:

5’-AGCGTGGGCGGGAAAATCTCTAGCA-3’
5’-ACTGCTAGAGATTTTCCCGCCCACGCT-3’

Sso7d-IN CSC intasomes were assembled and purified similar as previously described (*15*). Briefly, 3.0 uM integrase and 1.0 uM viral DNA substrate were mixed in the buffer containing 20 mM HEPES pH 7.5, 25% (w/v) glycerol, 50 mM NDSB-256, 5 mM b-ME, 4 μM ZnCl_2_, 100 mM NaCl, 5 mM CaCl_2_, or 5 mM MgCl_2_ with 50 uM dolutegravir (DTG) or 25 uM 4d. The reaction was carried at 37°C for 60 min and stopped by transferring onto ice, then, 500 mM NaCl was added to assembly reaction mixtures to solubilize intasomes. 10 uM of LEDGF-IBD was added into supernatant of intasomes, and incubated at 4°C for overnight. The reaction mix was loaded onto a HisTrap column (Cytiva) equilibrated with 20 mM Tris-Cl pH 8.0, 5mM 2-mercaptoethanol, 0.5 M NaCl, 20% (w/v) Glycerol, the intasomes were then eluted with a liner gradient of 0 mM to 500 mM imidazole in the same buffer. The fractions containing intasomes were combined and concentrated using a concentrator (Ultracel-100K, Millipore) and loaded onto Superose 6 10/300 GL (Cytiva) gel filtration column equilibrated with 20 mM Bis-Tris pH 6.2, 0.5 mM TCEP, 600 mM NaCl, 5.0 mM MgCl_2_, and 10% (w/v) glycerol. The intasomes were further purified with TSKgel UltraSW in the buffer of 20 mM Bis-Tris pH 6.2, 0.5 mM TCEP, 550 mM NaCl, 5.0 mM MgCl_2_, and 6% (w/v) glycerol. 25uM INSTI was added to the intasomes after every step of purification. CSCs were concentrated to 0.4 mg/ml with spin concentrator for cryo-EM grids preparation. Apo CSCs were prepared in essentially the same way except MgCl_2_ and INSTI were substituted by 5.0 mM CaCl_2_, and MgCl_2_ was omitted in all purification buffers.

### IN activity assays

3 uM integrase (unless otherwise noted) and 1.0 uM viral DNA substrate were preincubated on ice in 20 mM HEPES pH 7.5, 25% glycerol, 10 mM DTT, 5 mM MgCl2, 4 μM ZnCl2, and 100 mM NaCl in a 20 μl reaction volume. 300 ng of target plasmid DNA pGEM-9zf was then added and the reaction was initiated by transfer to 37°C and incubation was continued for 1 hr. For integration product analysis, the reactions were stopped by addition of SDS and EDTA to 0.2% and 10 mM, respectively, together with 5 μg of proteinase K. Incubation was continued at 37°C for a further 1 hr. The DNA was then recovered by ethanol precipitation and subjected to electrophoresis in a 1.5% agarose gel in 1x TBE buffer. DNA was visualized either by ethidium bromide staining or by fluorescence using a Typhoon 8600 fluorescence scanner (GE Healthcare).

### Modeling of double mutant cycles using Potts statistical energy models

The Potts model is a probabilistic model of sequence co-variation built on the single and pairwise site amino-acid frequencies of a protein multiple sequence alignment (MSA) and is aimed to describe the observation probabilities of the various states of the system (i.e., sequences in the MSA). The Potts statistical energy of a sequence provides an estimate of the probability of observing the sequence, giving a proxy for the fitness of the sequence, and for a sequence S of length L is given by 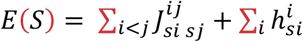, where the model parameters 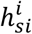 called ‘fields’ represent the statistical energy of a residue S_i_ at position i in sequence S, and 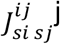 are ‘couplings’ representing the energy contribution of a position pair i,j. The probability of observing the sequence S according to the model is given by *P*(*S*) ∝ *e*^-*E*(*S*)^. The change in Potts energy due to mutating a residue a at position i in S to β is then given by:

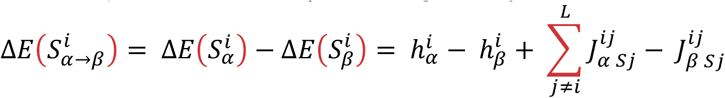

In this form, 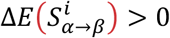 implies that the residue β is more favorable than residue a at position i in the given sequence S. For the pair of mutations α and β at positions i,j in the protein, the double mutant cycle captures the non-additivity (strength of epistatic interaction) of the effects of mutations and is given by the difference between the sum of the independent mutational effects, 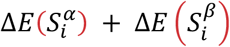, and the effect of the corresponding double mutation 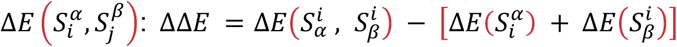. If ΔΔE=0, then the mutations are mutually independent, whereas if ΔΔE≠0, then the two mutations are epistatically coupled; ΔΔE>0 implies that the mutations are beneficial/co-operative to each other and *vice versa*.

The Potts statistical energy models probe the underlying relationships between protein structure, function, and fitness and, here, they are used to provide insights into the coupled nature of the effects of primary and compensatory residue pairs (see **Supplementary Note 1**). For a more detailed description of the Potts model in protein biophysics, we refer the reader to references (*52–56*), and for applications of the Potts model to HIV including description of data processing and model inference to references (*38, 42, 43, 57–60*).

### Cryo-EM sample preparation and data collection

For cryo-EM sample preparation, 3.0 ul of purified CSC was applied onto freshly plasma cleaned (60s, Pelco easyGlow plasma cleaner) holey grids (UltrAuFoil R 1.2/1.3 300 mesh, quantifoil), adsorbed for 10 sec, blotted for 4 seconds, then plunged into liquid ethane using Vitrobot V plunge freezer (Thermo Fisher). Data collections were performed at different locations using a Titan Krios transmission electron microscope (Thermo Fisher Scientific) operating at 300 keV, equipped with either a K2 or K3 direct electron detector (Gatan). Data collection was performed either using Leginon (*61, 62*) or SerialEM (*63*), depending on the facility being used. Most of the data collections were performed using stage tilt, respectively. All imaging parameters across the 11 datasets are comprehensively summarized in Table S1.

### Cryo-EM image analysis

Cryo-EM datasets were processed using a similar workflow. The movie frames were imported to Relion 4.0-beta-2 (*64*) for dose-weighted motion correction (*65*) on 5 by 5 patch squares using a B-factor of 150 Å^2^. The gain reference used for Relion motion correction was generated by using the Sum_all_tifs program, which is part of the cisTEM image processing suite (*66*). The motion-corrected micrographs were then imported into Warp 1.0.9 (*67*) to perform CTF estimation and particle selection. The particles that scored above 0.6 were selected in Warp with a re-trained BoxNet model using constraint settings of particle diameter of 180 Å and a minimum distance between particles of 20 Å. Particles were initially extracted in Warp using a boxsize of 384 pixels. The motion corrected micrographs in Relion and particle star file generated in Warp were then imported into cryoSPARC V3.3.2 (*68*) to continue data processing. The particle coordinates indicated in the particle star file were re-extracted in cryoSPARC with the same boxsize of 384 pixels and were used to perform 2D classification. The classes containing the best particles were selected after several rounds of 2D classification based on particle features and visual inspection of the 2D class averages. Following 2D classification and selection of the best classes, the remaining particles were subjected to heterogeneous refinement with C2 symmetry to further clean the dataset and using a previously published map of the HIV-1 intasome bound to the INSTI **4d** as a reference (EMD_20484) (*15*). Multiple rounds of heterogeneous refinement were performed until no further improvements in resolution were observed, as determined by the Fourier shell correlation (FSC). At each round, particles corresponding to the high-resolution map were selected, whereas the remaining particles were removed from processing. Following heterogeneous refinement, the remaining particles were used for homogeneous refinement in cryoSPARC, which generally produced reconstructions resolved to FSC criterion (*69*) using a fixed threshold of 0.143. At this point, the remaining particles were imported back to Relion for Bayesian polishing using default parameters. Subsequently, the polished particles were re-imported back to cryoSPARC to perform one round of non-uniform refinement (*70*), which was followed by one round of global CTF refinement and one round of perparticle CTF refinement. We repeated the procedure iteratively – Bayesian polishing in Relion followed by non-uniform refinement and global and per-particle CTF refinement in cryoSPARC iteratively, until no further improvements in resolution were observed, as determined using the FSC. Generally, the optics parameters, per-particle defoci, and the map resolutions converged after several iterations of polishing and non-uniform and CTF refinement. The final map was generated by the last non-uniform refinement. The directional resolution of the map was evaluated using the 3D FSC server (3dfsc.salk.edu) (*71*), and the quality of the orientation distribution was evaluated using the Sampling Compensation Function (SCF) (*72, 73*). All images were generated using UCSF Chimera (*74*).

### Atomic model building and refinement

Modeling of HIV-1 intasomes containing drug resistant variants was initialy focused on deriving a single accurate unhydrated atomic model. The initial reference used to derive an accurate model of the nucleoprotein components was the structure of the cleaved synaptic complex intasome bound with magnesium and the INSTI 4d, resolved to 2.8 Å (PDB 6PUY). A baseline atomic model was derived for the structure of the G140A intasome bound to DTG, which corresponded to one of the highest resolution maps, at 2.2 Å. Mutations corresponding to the drug resistant variants were generated in silico, the coordinates of the ligand (DTG) were added to the model, restraint files were generated using phenix.elbow (*75*), and the model was manually adjusted in Coot (*76, 77*) wherever discrepancies were apparent with the high-resolution map. The final model was derived through iterative model building using Coot and real-space refinement using phenix.real_space_refine (*78*). The same procedure was then adapted to all other map/model combinations, using the model of the G140A intasome bound to DTG as a baseline.

Atomic models were individually hydrated to account for variances in their global resolution. We determined an estimate for the number of waters that we should observe based on an analysis of Carugo and Bordo (*79*). Before dousing, cryo-EM density reconstructions were resampled and resized such that the voxel size became roughly one quarter of the resolution estimate using software available in CisTEM (*66*). We next used phenix.douse, available from Phenix version 1.20.1-4487 (*80*) to programmatically add water molecules to the density that was not accounted by the protein or DNA models. During dousing, we used the same settings for each of the models – dist_min=2.5 dist_max=4.5 keep_input_water=TRUE sphericity_filter=false – and varied the map_threshold value to add the number of waters expected based on the resolution of the map and the number of atoms in the model according to analysis of Carugo and Bordo. After dousing, the models were subjected to a round of real space refinement with phenix.real_space_refine. Each map was then manually inspected to evaluate agreement of water placement with cryo-EM densities. Problematic water molecules were identified using validation tools in Phenix (*81*) and Molprobity (UnDowser) (*82*) and removed if they (1) clashed with protein molecules, (2) were placed in density that belong to other protomers of the dodecamer, or (3) were placed errantly without corresponding density. Dousing was performed using an asymmetric unit of the intasome, and two-fold symmetry was applied at the end. The geometry of the final symmetrized models and other validation statistics were reported by Molprobity. The relevant refinement statistics are summarized in **Table S1**. High-resolution structural figures were prepared for publication using UCSF Chimera (*74*).

### Molecular dynamics and free energy perturbations system setup and simulation details

For MD-based FEP simulations, the starting model was an experimentally determined structure of the E138K/Q148K HIV intasome with **DTG** bound. The structure of the E138K/Q148K-**4d** complex used in the FEP simulation is obtained from E138K/Q148K-**DTG**, but replacing **DTG** with **4d** from the WT intasome with **4d** bound (PDB 6PUY) (*15*)). The E138K/Q148K-DTG and E138K/Q148K-**4d** complexes were prepared with Protein Preparation Wizard. FEP simulations using the FEP+ program (*83*) from the Schrödinger suite 2021-2 was employed to mutate Gly140 into Ala. A series of FEP simulations are performed to alchemically “mutate” the Gly140 side chain into Ala. The structure of the complex E138K/G140A/Q148K-ligand corresponds to the final state (< = 1) of the FEP simulation. For every complex and the corresponding endpoint structures, the FEP Protein Mutation for ligand Selectivity GUI (*83*) in Maestro Suite is applied to build a perturbation map. The OPLS4e force field was used for modeling proteins and ligands (*84*). Torsion parameters were checked for all ligand fragments using Force Field Builder. A 10 Å cubic box filled with ~31,300 SPC water (*85*) was used for the complex and solvent perturbation leg. Additional Na+ and Cl-ions were added to achieve 0.15 M NaCl in the simulation box. The number of alchemical λ windows is set to 12 by default to connect the wild-type and the mutant states. For each λ window, the production MD is run for 15ns in the NPT ensemble. The Bennett acceptance ratio method (*86*) (BAR) was used to calculate the free energy differences. The mutational free energy calculations were performed in triplicate by initializing the MD with different random seeds. The reported relative binding free energies were calculated as the average of the three ΔΔG from independent simulations.

## SUPPLEMENTARY MATERIAL

### Supplementary Text

#### Supplementary Note 1: Modeling the fitness landscape using Potts statistical energy models built on the Stanford drug resistance DB

A hallmark of drug resistance is the pairing of one or more primary mutations that interfere with drug binding and for which there is often a fitness cost, with one or more compensatory mutations that improves the fitness in the presence of the primary mutation(s). The association of primary with compensatory mutations is a feature of epistasis, *i.e*. the coupling of fitness effects that are not independent (*43*). The Potts statistical energy models probe the underlying relationships between protein structure, function, and fitness and, here, they provide insight into coupled primary and compensatory residue pairs that help to explain the diverse data at hand.

The population-based Potts model of the fitness landscape is defined though an expression which defines the likelihood of a sequence in a given environment, and it changes as the environment changes. It has the following functional form: 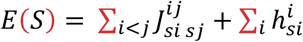 where the residue-residue couplings 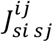 correspond to epistatic interactions involving compensatory constraints between residues within the protein, and the field terms 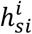 represent single-site biases. A parsimonious interpretation of these two terms is that the compensatory constraints encoded in the couplings, like those that enforce correct folding, are “intrinsic” and operate in all the environments, while the field terms correspond to “external” selection pressures, like the immune response and drug selection pressure. Fitness is always defined relative to an environment, and in this decomposition of the fitness, different environments (for instance different drug regimens) will correspond to different parameterizations of only the field terms.

Our Potts model is fit to drug-exposed sequences (mainly exposed to **RAL**, with a minority exposed to **EVG**, **CAB**, **DTG**, and **BIC**), and therefore models a global drug-experienced fitness environment for the virus, reflecting its survival as it evolves across hosts given combinations of these drugs. As anticipated by our interpretation of the physical meaning of the fields and couplings presented above, we have confirmed that starting with the drug experienced model, we can transform it to model the drug-naïve fitness landscape which accurately predicts mutant frequencies of a drug-naïve sequence dataset by modifying only the field terms of the drug experienced model. Both this drug-experienced model and the drug-naive model predict the respective pairwise mutant frequencies in each dataset with a maximum of 1% error, and on average only 0.07% error, even though the pairwise mutant frequencies of the drug-naïve dataset are up to 53% different from the drug-experienced mutant frequencies. These two models share the same (intrinsic) coupling terms, but have different (external) field terms.

Double-mutant-cycles, defined as ΔΔ*E* in Methods, equal the fitness effect of the double mutant minus the sum of the two single mutant fitness effects; the external field terms of the Potts model cancel out of this computation. Therefore, the double-mutant-cycle values have the special property that they reflect epistatic constraints within the enzyme present in all environments, and their values are the same for both the drug-experienced and derived drug-naïve model described above. This provides the rationale for our using the predictions of double-mutant-cycles to estimate intrinsic epistatic effects that are independent of drug exposure, even though our Potts model is fit to data exposed mainly to **RAL**. We perform our inference and computations using the drug-experienced dataset, because it has more extensive sequence variation than the drug-naïve data, giving a more accurate model parameterization of the intrinsic epistatic interactions.

The results of the double mutant cycle analysis (**Fig. 2D**) show the non-additive effects (cooperativity) between compensatory pairs of mutations in the protein sequence. The double mutant cycle values DDE depend only on the Potts model coupling terms l(i_a_,j_b_), not the field terms. The coupled mutation pairs involving residues at positions 138, 140, and 148, which are the central focus of the current work, are among those with the strongest epistatic couplings. While the majority of drug experienced data present in the Stanford DB are derived from patients who are treated with 1^st^ generation INSTIs, the couplings derived from this dataset reflect epistatic effects due to intrinsic constraints on fitness induced by the structure and function of the enzyme, independent of the application of a specific drug. This reinforces the view that the epistatic interactions among mutations at positions (138,140,148) that are captured by the Potts model reflect fitness constraints imposed by the structure and function of the protein, independent of a specific drug.

#### Supplementary Note 2. Correlation between FEP-based binding affinities and experimental measurements of EC_50_ in cells

We show that experimental EC_50_ values correspond closely to the calculated dissociation constant K_D_. As seen in **table S2**, based on the FEP calculations, we estimate that with the introduction of the third mutation, G140A, the binding affinities of **DTG** and **4d** are reduced by 1.0 and 0.3 kcal/mol (for dominant conformers), respectively, which correlates well with the experimentally determined EC_50_ values from virology assays. This result suggests that the atomistic free energy model captures the trend of the drug resistance mutation of the IN to the INSTIs, and such techniques may further improve our understanding of the drug resistance mutations to the INSTIs.

#### Supplementary Note 3. 4d binds more strongly to the E138K/G140A/Q148K triple mutant when the adenine base forms a stacking interaction with the ligand

A comparison of the Apo structure (PDB: 6PUT) (*15*) to the INSTI-bound structures of the Intasome shows that the difference is largely confined to the conformation of the terminal adenosine (dA21) of the vDNA, which can adopt two distinct conformations in the active site of INSTI-bound structures: (i) one in which the dA21 base forms a stacking interaction with the bound INSTI (stacked) and (ii) one in which the base extends into the inter-subunit space (non-stacked). Both structures are distinct from the conformation of vDNA in the apo cryo-EM structure. Using the thermodynamic cycle shown in **Fig. 5G**, we can gain insight into the relative binding affinities for the ligands **DTG** and **4d** to the stacked and non-stacked conformations using as constraints the experimental EC_50_ values (estimate of the dissociation constant K_D_) and stacking vs. non-stacking ratios estimated from the cryo-EM densities. There are two constraint equations, which can be used to solve for the binding strength of **DTG** or **4d** to the stacked form of the complex Δ*G^S^* and to the non-stacked form of the complex Δ*G^NS^*. According to the thermodynamic cycle shown in **fig. 5G**, 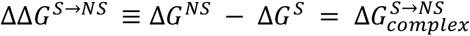, which means that we can estimate the difference in the ligand binding free energy towards the stacked vs. non-stacked conformations ΔΔ*G*^*S*→*NS*^ = Δ*G^NS^* - Δ*G^S^* from the knowledge of 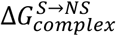. Because 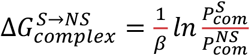, the ratio of the occupancies of the stacked vs. non-stacked complex states 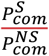 can be obtained from experiments. **Table S3** summarizes the experimental constraints and shows that **4d** binds to KAK in both non-stacked and stacked conformers more strongly than **DTG** from the estimated binding free energies. Furthermore, the binding free energy difference between **4d** and **DTG** to E138K/G140A/Q148K is more negative (−2.1 kcal/mol) in the stacked conformer than in the non-stacked conformer (−0.9 kcal/mol), indicating that the stacked terminal adenine in E138K/G140A/Q148K helps maintain bound **4d** in its preferred position, and contributes more to the greater potency of **4d** than does the non-stacked conformation.

### Supplementary Figures

**Figure S1:**
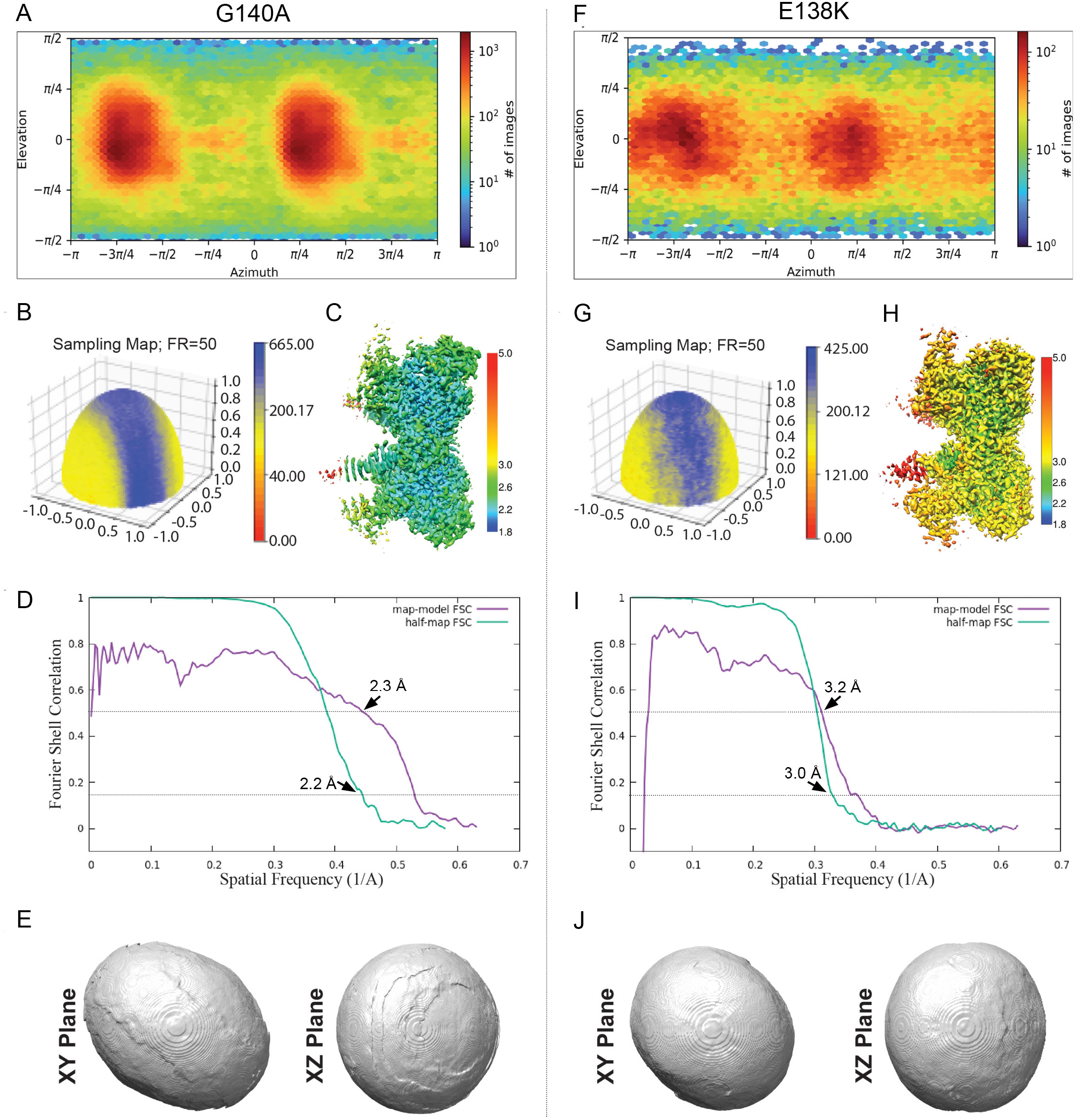
Cryo-EM validation of the highest resolution G140-DTG and the lowest resolution E138K-DTG intasome reconstructions. Validations metrics are displayed for (**A-E**) the G140A intasome with DTG bound or (**F-J**) the E138K intasome with DTG bound. (**A,F**) Euler angle plot showing the distribution of projection orientations used in the cryo-EM reconstruction. (**B,G**) Surface sampling plot of the Fourier voxel sampling derived from the Euler angle distribution plot, calculated with a Fourier radius set to 50 voxels and imposing C2 symmetry. The sampling compensation factor (SCF) is indicated (*72, 73*). (**C,H**) Cryo-EM reconstruction colored by local resolution. (**D,I**) Fourier shell correlation (FSC) curves derived from half-map and map-to-model reconstructions, with FSC cutoffs 0.143 and 0.5 indicated, respectively, as well as nominal resolution values. (**E**,**J**) 3DFSC (*71, 87*) shown as an isosurface and thresholded using a value of 0.5 with two perpendicular planar views.

**Figure S2:**
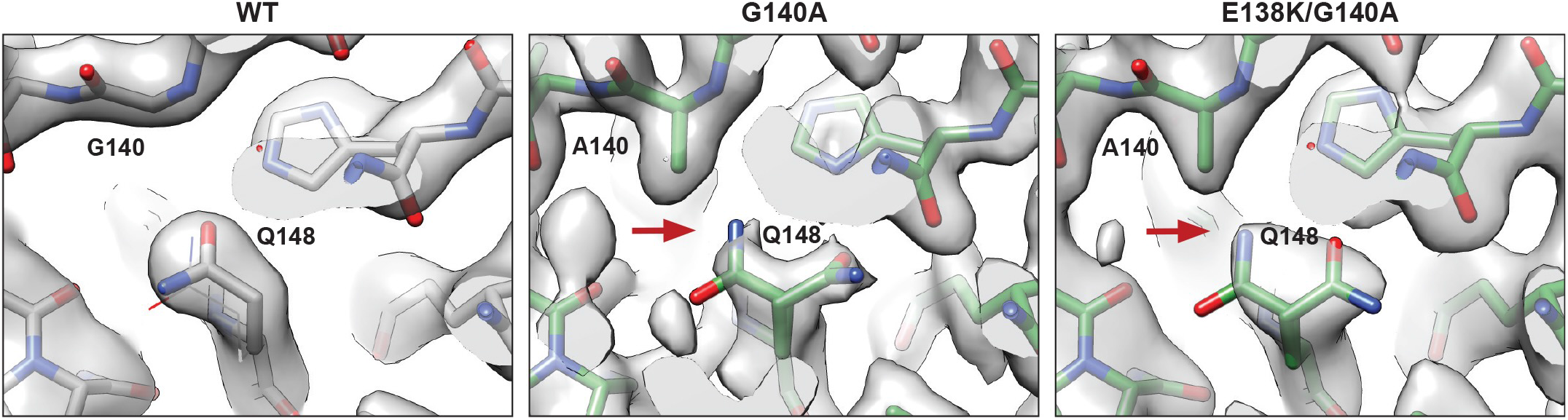
Rotameric configurations of Q148 in the presence of the G140A mutation and the E138K/G140 Double mutation. Cryo-EM density and the atomic models are shown for structures of WT, G140A, and E138K/G140A intasomes bound to **DTG**. Only one rotamer of Q148 is evident for the WT intasome, whereas two rotameric conformations can be modeled for G140A and E138K/G140A mutant intasomes. There is a preference for the alternative conformation of Q148 due to weak steric clashes (indicated by red arrows) if A140 is present. Residues at positions 140 and 148 are labeled.

**Figure S3:**
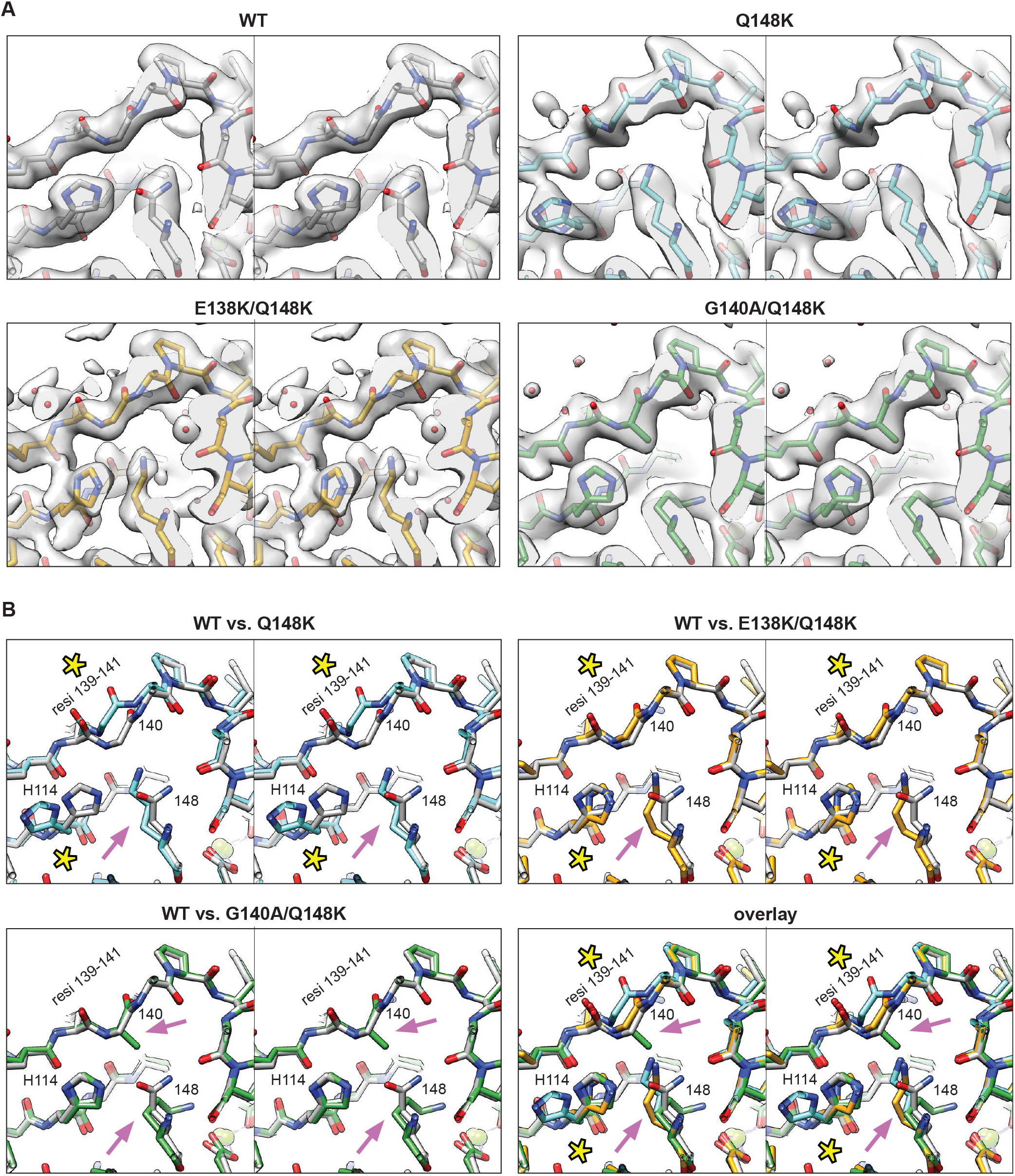
Changes to backbone residues 139-141 and the sidechain of H114 if the Q148K mutation is present. (**A**) Experimental cryo-EM density and atomic models of the region surrounding residue 148 are shown in stereo for structures of WT, Q148K, E138K/Q148K, and G140A/Q148K intasomes bound to **DTG**. (**B**). Overlays of atomic models showing the same region displayed in panel A. In comparison to WT, the backbone shift is readily apparent in the Q148K structure, and there is a small shift in the E138K/Q148K structure. The reconfiguration of the rotameric position of H114 is apparent in Q148K structure, whereas only a minor rotation is apparent in E138K/Q148K structure. Residues at positions 114 and 148 are labeled.

**Figure S4:**
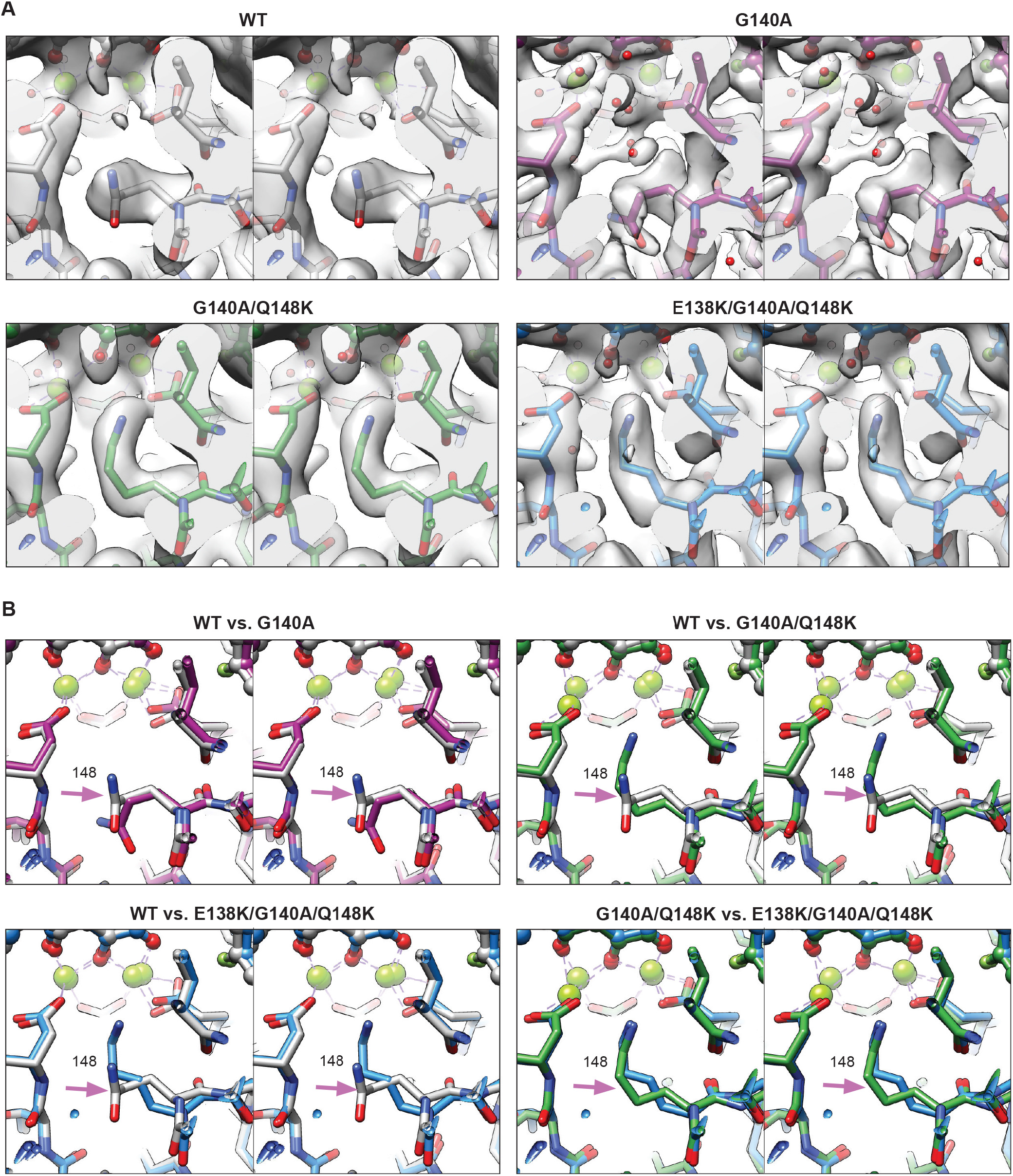
Alterations in the conformation of the residue at position 148 in response to the G140A mutation. (**A**) Experimental cryo-EM density and the atomic models centered around residue 148 are shown in stereo for structures of WT, G140A, E138K/Q148K, and G140A/Q148K intasomes bound to **DTG**. (**B**). Overlays of atomic models, showing the same region displayed as in panel A. In comparison to WT, the conformational change of Q148 is apparent if G140A is present. Furthermore, if the residue at position 148 is a Lys, the presence of the G140A mutation leads to a reconfiguration of the Lys sidechain to position that is closer to the Mg^2+^ ions, as shown in the structures of G140A/Q148K and E138K/G140A/Q148K. Position 148 is labeled. Red arrows indicate weak steric clashes.

**Figure S5:**
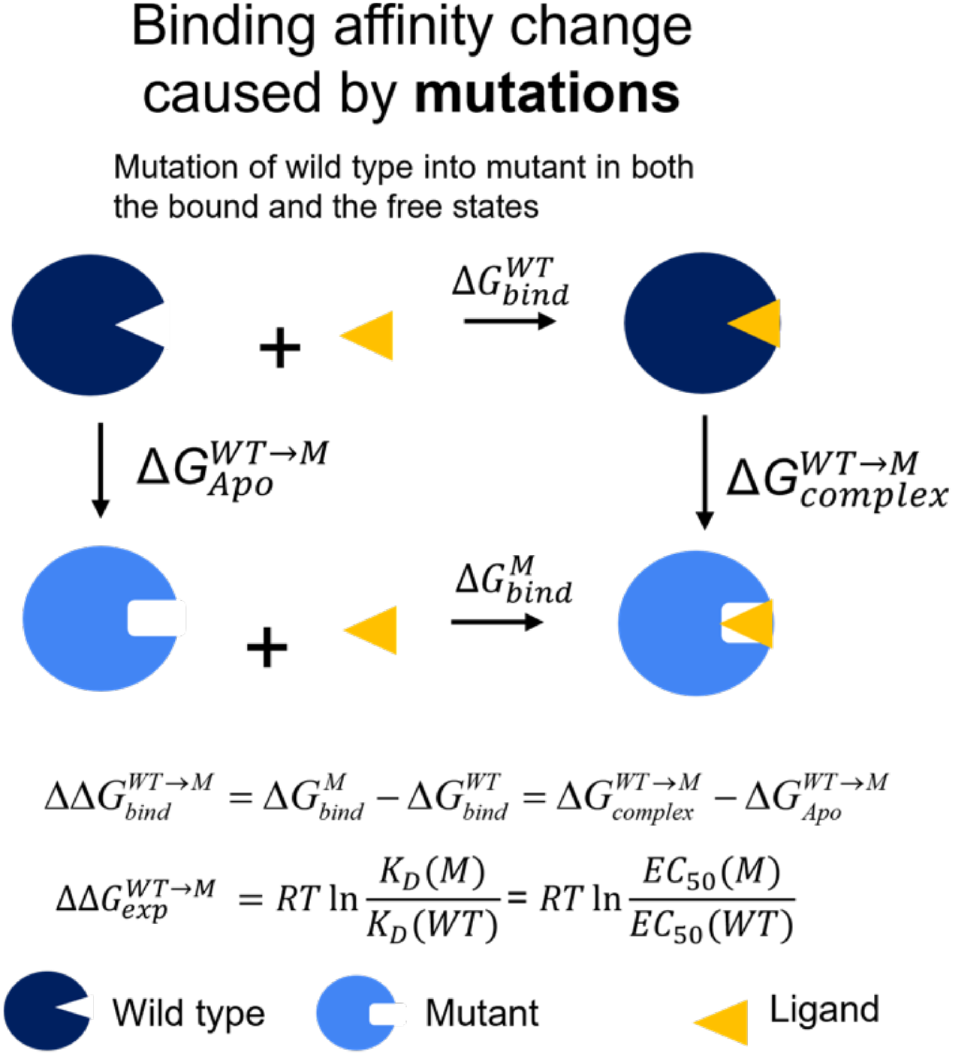
Thermodynamic cycles used for MD-based FEP simulations. The thermodynamic cycle used for computing the changes in the binding free energy caused by the residue mutation in the binding of the ligand to the intasome. We examined the changes in the binding free energy caused by the G140A mutation in the binding of either **DTG** or **4d** to either the E138K/Q148K (KK) or E138K/G140A/Q148K (KAK) DRM intasomes by using FEP (free energy perturbation) to calculate the relative binding free energy 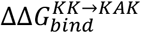 for each of the ligands. In the thermodynamic cycle, 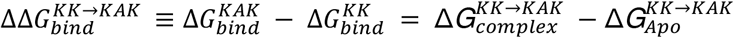, i.e. the relative binding free energy 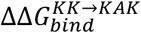 can be obtained by the two vertical legs, instead of simulating the physical binding process.

**Figure S6:**
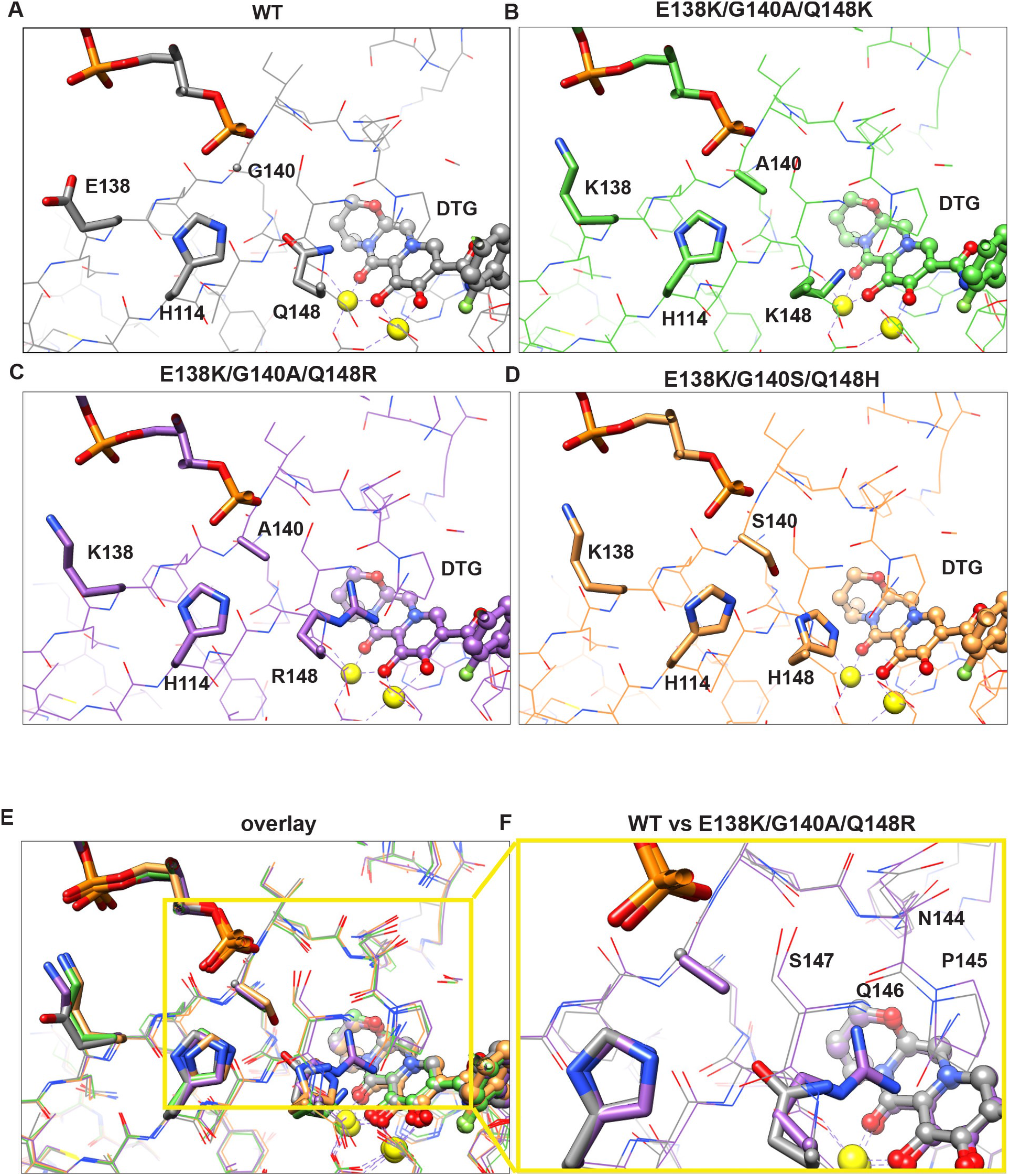
Comparison of the structural changes in response to Lys, Arg, and His mutations at position 148. Atomic models of the active site from high-resolution intasome structures with **DTG** bound are shown for (**A**) the WT intasome, or (**B-D**) the triple mutant variants (**B**) E138K/G140A/Q148K (**C**) E138K/G140A/Q148R, (**D**) E138K/G140S/Q148H. (**E**) An overlay of all structures in A-D. (**F**) Close-up of the region spanning N144-S147, which differs in the structure of E138K/G140A/Q148R intasome with **DTG** bound, compared with the WT intasome with **DTG** bound. The view of the active site is the same as in Figure 3.

**Figure S7:**
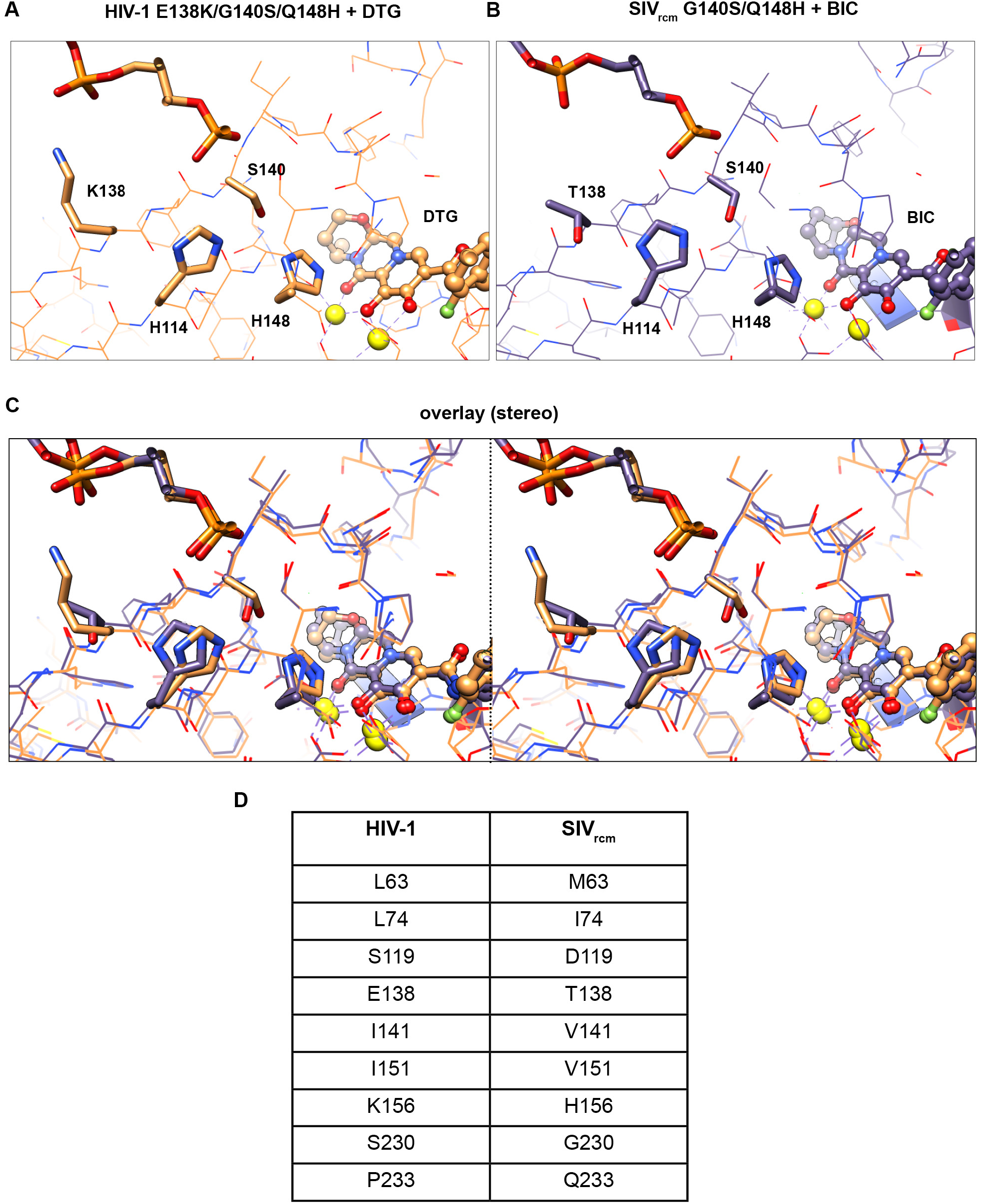
Comparison between the structures of HIV-1 and SIVrcm intasomes containing the G140S/Q148H mutations. Atomic models of the active site from high-resolution intasome structures are displayed for (**A**) the HIV-1 E138K/G140S/Q148H intasome with **DTG** bound, and (**B**) the SIV_rcm_ G140S/Q148H intasome with BIC bound. (**C**) An overlay of the two structures in A-B, displayed in wall-eye stereo. (**D**) A table of the distinct residues within 10 Å of the bound drugs which differ between HIV-1 and SIV_rcm_ IN. In **panels A-C**, the view of the active site is the same as in **Figure 3**.

**Figure S8:**
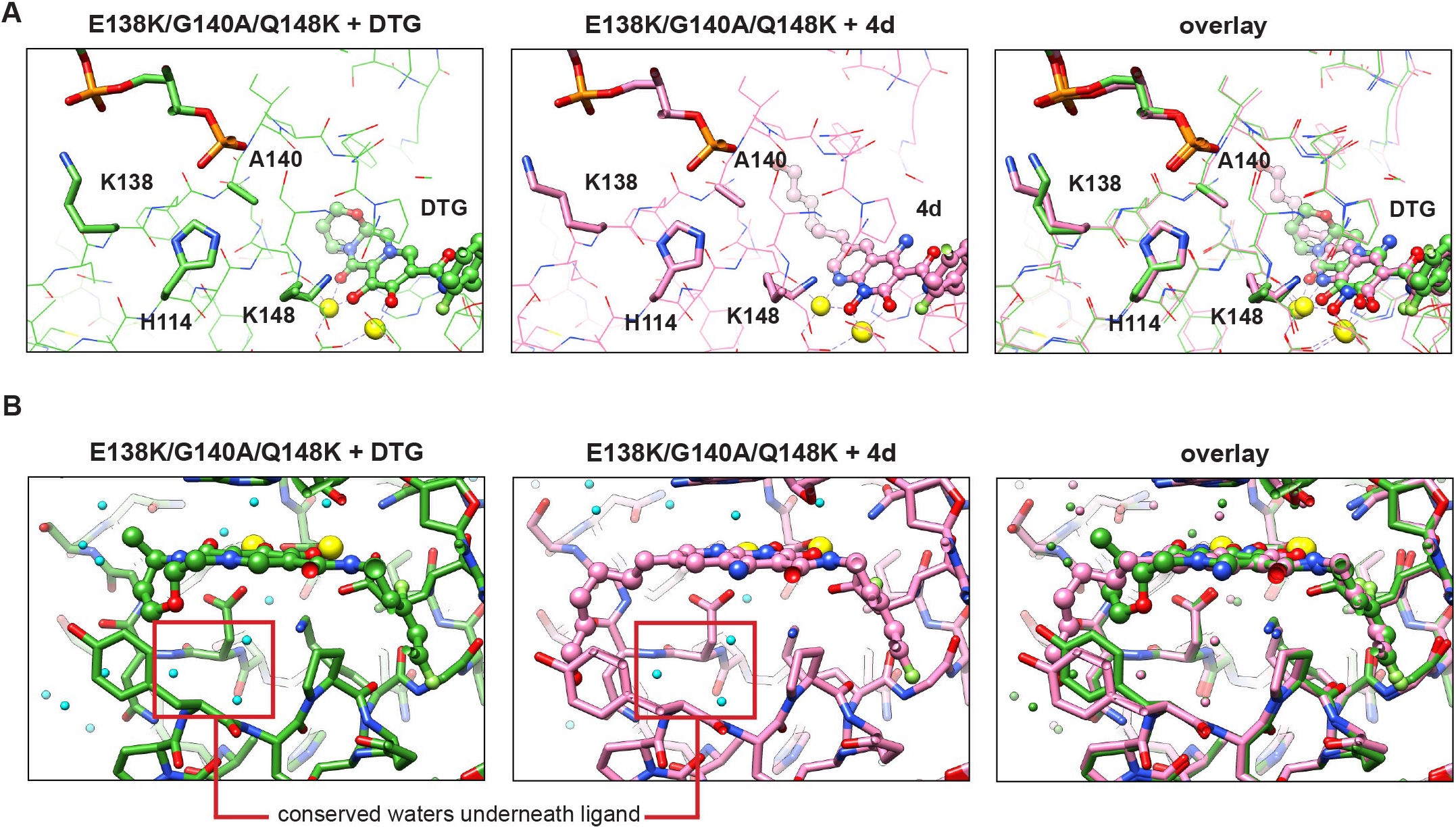
Comparison of the structural changes in the intasome when 4d or DTG are bound. (**A**) Atomic models of the active site from high-resolution HIV-1 intasome structures of the E138K/G140A/Q148K variant bound to either **DTG** or **4d**, with an overlay of the two at right. Only subtle changes are observed for the residues involved in the resistance mechanism. The view is identical to that in Figure 3. (**B**) Atomic models of the region surrounding the bound ligand, showing bound water molecules for each structure (cyan). An overlay of the two is at right, and the waters are colored according to its respective structure. The waters outlines in red, which reside “underneath” the ligand, are entirely conserved for the two mutants. Minor variations in the hydration shell is observed otherwise, which may arise from biological differences as well as differences in resolution.

**Figure S9:**
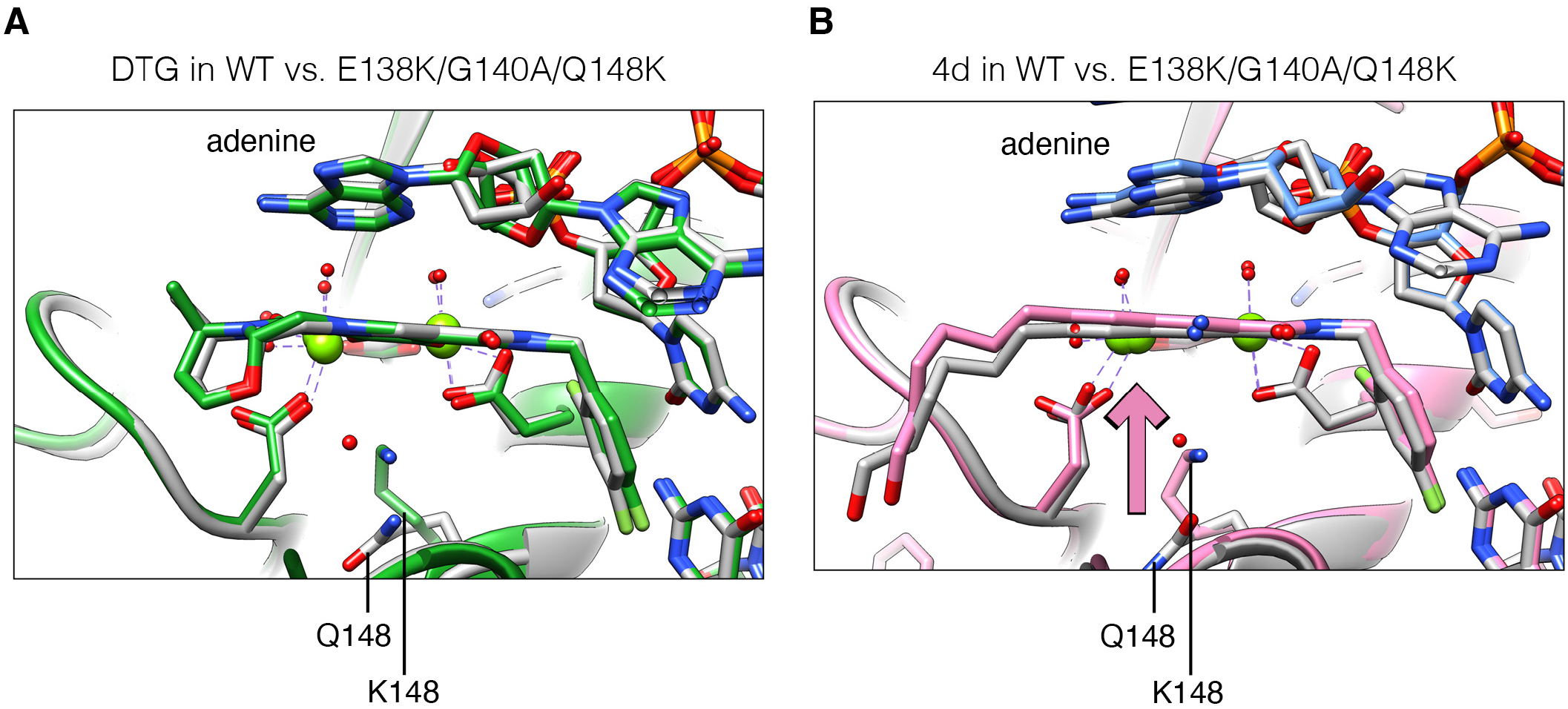
The ligand 4d, but not DTG, is pushed toward the terminal adenine base in the E138K/G140A/Q148K intasome. (**A**) Overlay of the atomic models of WT (gray) and E138K/G140A/Q148K (green) intasome with **DTG** bound. The ligand remains in place in the context of the triple DRM. (**B**) Overlay of the atomic models of WT (gray) and E138K/G140A/Q148K (pink) intasome with **4d** bound. The ligand is pushed “upward” in the context of the triple DRM (pink arrow).

**Table S1.** Cryo-EM data collection and image processing parameters. See excel sheet attached

**Table S2.** Changes in the ligand binding free energies caused by the G140A mutation (E138K/Q148K à E138K/G140A/Q148K) for the INSTIS **DTG** and **4d** bound to IN. The first column shows the binding free energy differences from EC_50_ values 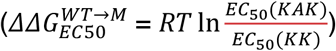 of **DTG** and **4d** against the mutant IN E138K/Q148K and E138K/G140A/Q148K estimated using FEP simulations, whereas the last column shows the experimental binding free energy differences obtained using virology assays (**Fig. 1**).

| ligand | FEP Calculated $\Delta\Delta G(\text{kcal/mol})$<br>$= \Delta G_{bind}^{KAK} - \Delta G_{bind}^{KK}$ | Experimental<br>$\Delta\Delta G(\text{kcal/mol})$<br>$= RT \ln \frac{EC_{50}(KAK)}{EC_{50}(KK)}$ |
| --- | --- | --- |
| DTG | 1.0 ± 0.6 | 1.3 |
| 4d | 0.3 ± 0.4 | 0.1 |

**Table S3.** The estimated binding free energies for **DTG** and **4d** bound to WT and E138K/G140A/Q148K intasomes in the stacked and non-stacked conformations.

|  | DTG-WT | 4d-WT | DTG-KAK | 4d-KAK | Notes |
| --- | --- | --- | --- | --- | --- |
| The probability of the complex in stacked conformer $P_{com}^S$ | 0.58 | 0.65 | 0.40 | 0.82 | Occupancies from cryo-EM structures. |
| The probability of the complex in non-stacked conformer $P_{com}^{NS}$ | 0.42 | 0.35 | 0.60 | 0.18 | |
| Binding free energy of ligand bound to IN in stacked conformer $\Delta G^S$ | -11.4 | -11.8 | -8.9 | -11.0 | Solved with the following two constraints |
| Binding free energy of ligand bound to IN in non-stacked conformer $\Delta G^{NS}$ | -11.2 | -11.4 | -9.2 | -10.1 | |
| overall binding free energy of ligand to IN<br>$\Delta G_{bind} = -\frac{1}{\beta} \ln (e^{-\beta\Delta G^S} + e^{-\beta\Delta G^{NS}})$ | -11.7 | -12.1 | -9.5 | -11.1 | Constraint 1. (Section 1)<br>$\Delta G_{bind} = \frac{1}{\beta} \ln (EC50)$ |
| EC50(nM) | 2.6 | 1.5 | 117.6 | 7.3 |  |
| Binding free energy difference $\Delta G^{NS} - \Delta G^S$ | 0.2 | 0.4 | -0.2 | 0.9 | Constraint 2. (Details in Fig 1)<br>$\frac{1}{\beta} \ln \frac{P_{com}^S}{P_{com}^{NS}}$ |
| $\Delta G^S_{4d} - \Delta G^S_{DTG}$ | | -0.4 | | -2.1 | |
| $\Delta G^{NS}_{4d} - \Delta G^{NS}_{DTG}$ | | -0.2 | | -0.9 | |

